# Nodule organogenesis in *Medicago truncatula* requires local stage-specific auxin biosynthesis and transport

**DOI:** 10.1101/2023.12.07.570677

**Authors:** TT Xiao, D Shen, S Müller, J Liu, A van Seters, H Franssen, T Bisseling, O Kulikova, W Kohlen

**Affiliations:** Department of Plant Sciences, Laboratory of Molecular Biology, Cluster of Plant Developmental Biology, Wageningen University, Droevendaalsesteeg 1, 6708 PB, Wageningen, The Netherlands; Current address, College of Plant Science and Technology, Beijing Key Laboratory for Agricultural Application and New Technique, Beijing University of Agriculture, Beijing 102206, China; Current address, Max Planck Institute for Plant Breeding Research, Carl-von-Linne-Weg 10, D-50829 Cologne, Germany; Department of Plant Sciences, Laboratory of Cell and Developmental Biology, Cluster of Plant Developmental Biology, Wageningen University, Droevendaalsesteeg 1, 6708 PB, Wageningen, The Netherlands

**Keywords:** Medicago, Rhizobium, root nodule development, RNA *in situ*, local auxin biosynthesis, auxin transport, DR5 patterns, auxin gradient

## Abstract

The importance of auxin in plant organ development including root nodule formation is well established. Using auxin reporter constructs the spatiotemporal auxin distribution pattern during nodule development has previously been illustrated. However, our understanding of how this pattern is built-up and maintained still remains elusive.

To this end, we studied how the auxin gradient visualized by DR5 expression patterns at different stages of nodule development in Medicago truncatula (Medicago), is correlated with the spatiotemporal expression patterns of known auxin biosynthesis and auxin transport genes. In addition, we record the MtPIN10-GFP expression pattern and polar positioning on the cell plasma membranes during nodule primordium development to investigate the auxin flux. RNA interference and the application of auxin synthesis blockers were used to demonstrate the relevance of biosynthesis and transport at the initial stages of the nodulation process.

Our results show that upon rhizobium inoculation, preceding the first mitotic activity, a specific set of MtYUCs and MtPINs as well as MtLAX2 are expressed in the pericycle contributing to the creation of an auxin maximum. Overall, we demonstrate that dynamic spatiotemporal expression of both, MtYUCs and MtPINs, result in specific auxin outputs in subsequent stages of nodule primordia and nodule meristem formation.

## INTRODUCTION

In legumes Rhizobium can establish a symbiotic relationship leading to the formation of nitrogen-fixing root nodules, involving two different developmental programs: infection thread initiation in the epidermis and nodule primordium formation in inner root cell layers. Nodule formation is initiated by lipo-oligosaccharides, called Nod factors, secreted by rhizobia (reviewed by Dénarié *et al*., 1996; and Geurts and Bisseling, 2002). In *Medicago truncatula* (Medicago), rhizobia enter the root via root hairs, where tubular structures, called infection threads, are formed (reviewed by Brewin, 2004; and Gage, 2004). Concomitantly, differentiated root cells, including pericycle and cortical cells are mitotically activated in an organized manner by which the Medicago nodule primordium is formed (reviewed by Oldroyd and Downie, 2006; Xiao *et al*., 2014). The infection thread that contains rhizobia, grows towards the primordium and bacteria are released in cells derived from the inner cortex. A meristem which facilitates further growth is formed at the apex of the nodule primordium.

In general, two types of nodules can be distinguished based on the lifespan of the meristem. In determinate nodules (e.g. *Lotus japonicus*, Lotus) the meristem is active for a short time, while in indeterminate nodules (e.g. Medicago) a persistent meristem is formed (reviewed by Hirsch, 1992; and Sprent and James, 2007). These plants also differ in the location of first cell divisions initiated upon rhizobial inoculation. Indeterminate nodules are initiated in the pericycle and the inner cortical cells and determinate nodules start in the outer and middle cortical cells (reviewed by Hirsch, 1992). In each nodule type the initiation of cell divisions is correlated with auxin accumulation (Mathesius *et al*., 1998*b*; Pacios-Bras *et al*., 2003; Wasson *et al*., 2006; Mathesius, 2008; Zhang *et al*., 2009; Ng *et al*., 2015; Herrbach *et al*., 2017; Ng and Mathesius, 2018).

Irrespective of the nodule type formed, auxin is involved in cell cycle control, vascular tissue differentiation and rhizobial infection during nodule development (reviewed by Kohlen *et al*., 2018; and Lin *et al*., 2020). Auxin is mainly synthetized in the aerial part of plant and transported along phloem passively and by auxin transporters actively into the root apex (Swarup *et al*., 2001; Vanneste and Friml, 2009; Swarup and Péret, 2012; Swarup and Bennett, 2014). Several studies have shown that inhibitors that block this shoot-rootward auxin transport induce the formation of pseudo-nodule by creating a local auxin maximum (Allen *et al*., 1953; Hirsch *et al*., 1989; Scheres *et al*., 1992; Schnabel and Frugoli, 2004; Subramanian *et al*., 2006; Wasson *et al*., 2006; Zhang *et al*., 2009; Rightmyer and Long, 2011; Shen *et al*., 2015; Sańko-Sawczenko *et al*., 2016). The inhibition of rootward auxin transport at the very beginning of nodule initiation was observed during initiation of indeterminate nodule formation but not for determinate nodules where, on the contrary, an increase in acropetal auxin transport was found (Allen *et al*., 1953; Hirsch *et al*., 1989; Rubery and Jacobs, 1990; Mathesius *et al*., 1998*b*; Boot *et al*., 1999; Pacios-Bras *et al*., 2003; Prayitno *et al*., 2006; Subramanian *et al*., 2006). Auxin polar transport is mediated by efflux carriers, among them the best studied are PIN-formed (PINs), and auxin influx carriers AUXIN RESISTANT1 (AUX1) and LIKE-AUX1 (LAX) (Petrásek *et al*., 2006; Swarup and Bhosale, 2019). PIN-dependent local auxin gradient is a part of all organ development processes in plants (Benková *et al*., 2003; Wisniewska *et al*., 2006). 12 *PIN* genes were identified in Medicago (Schnabel and Frugoli, 2004; Shen *et al*., 2015; Sańko-Sawczenko *et al*., 2016; Kohlen *et al*., 2018). All Medicago *PIN* genes have an orthologue in Arabidopsis (Schnabel and Frugoli, 2004; Peng *et al*., 2013; Sańko-Sawczenko *et al*., 2016; Kohlen *et al*., 2018). The expression pattern of *MtPINs* during nodule initiation and development, and reported phenotypes of *MtPIN*-RNAi plants in Medicago indicate their important role for the auxin transport during indeterminate nodule development (Huo *et al*., 2006; Shen *et al*., 2015; Sańko-Sawczenko *et al*., 2016; Schiessl *et al*., 2019).

Next to PINs, AUX1/LAX proteins function as auxin influx carriers. It has been reported that in Arabidopsis AUX1 and LAX3 play key roles during primary and lateral root development (Bennett *et al*., 1996; Marchant *et al*., 1999; Swarup *et al*., 2008). There are five members of the AUX1/LAX family in Medicago (Schnabel and Frugoli, 2004). The expression of *MtLAX2*, a homologue of *AtAUX1*, is highly induced during nodule primordia initiation (Roy *et al*., 2017; Schiessl *et al*., 2019). Moreover, two identified Medicago *Tnt1* insertion mutants effected in this gene, *Mtlax2-1 and Mtlax2-2*, developed fewer nodules and lateral roots, suggesting the same requirements for auxin influx activity for development of both lateral organs (de Billy *et al*., 2001; Schnabel and Frugoli, 2004; Swarup and Péret, 2012; Roy *et al*., 2017). In line with the putative function of the MtLAX2 protein, both mutants displayed decreased DR5-GUS activity associated with rhizobial infection (Roy *et al*., 2017).

Auxin maxima are not only established as the result of polar auxin transport (PAT) but also as a consequence of local auxin biosynthesis (reviewed by Zhao, 2018; Cao *et al*., 2019). Although there are several pathways that lead to the production of auxin, the TAA/YUC pathway is reported to be the primary endogenous auxin biosynthesis route (reviewed by Zhao, 2010, 2018). The TAA/YUC pathway is a two-step pathway in which TAA (TRYPTOPHAN AMINOTRANSFERASE) first converts tryptophane into indole-3-pyruvic acid (IPyA). Next, YUCCA, a flavin-containing monooxygenase, catalyses the rate-limiting step in synthesis of indole-3-acetic acid (IAA), the main natural auxin, from indole-3-pyruvic acid (IPyA) (Mashiguchi *et al*., 2011; Stepanova *et al*., 2011; Won *et al*., 2011). In Medicago four *YUCCA* (*YUC*) genes were reported to be upregulated during nodule primordium formation: *MtYUC1*, *MtYUC*2*, MtYUC8* and *MtYUC9* and their expression is (in)directly regulated by the root nodule symbiosis specific transcription factor NIN (Liu *et al*., 2019; Schiessl *et al*., 2019).

Overall, during nodule initiation, it is likely that defined auxin concentrations are needed to induce and guide nodule organogenesis. However, how the auxin related gene network is contributing to the establishment of this remains unknown. To unravel the complexities of these highly interconnected molecular physiological networks, we have employed a combination of techniques, including *DR5::GUS* based visualization of auxin output and the spatiotemporal expression profiling of genes associated with auxin synthesis and transport, known to be expressed during the formation of a nodule primordium. We observed that the accumulation of auxin precedes cell divisions and, in subsequent stages of Medicago primordium development, remains closely associated with mitotic re-activation. Our investigations have revealed both, distinct and overlapping spatiotemporal expression patterns for *MtYUCs* and *MtPINs* genes. Additionally, we have pinpointed the localization of *Mt*PIN10 at the inception of nodule initiation and during subsequent developmental stages. Furthermore, through the utilization of *RNAi* and pharmacological interventions, we have validated that in addition to auxin transport, local auxin biosynthesis is an indispensable contributor to the proper initiation and development of root nodules in Medicago.

## RESULTS

### Defining the auxin response pattern during nodule initiation

To create a frame of reference for spatiotemporal analyses of genes involved in auxin patterning, we first determined the dynamics of the auxin gradient during nodule primordium initiation and development as previously described (Xiao *et al*., 2014). For this, we created a stable transgenic Medicago R108 line containing the synthetic auxin-response promoter DR5 fused to β-glucuronidase (*DR5::GUS*).

Roots of these plants were collected at different time points after spot-inoculation with *Sinorhizobium meliloti* 2011 *(S. meliloti* 2011), and incubated in GUS substrate buffer. Segments of stained roots were cut, fixed, embedded in plastic and subsequently sectioned (Fig. 1). The earliest *DR5* activity was observed in pericycle cells prior to periclinal divisions. As this stage is not visible when scored for cytological changes as a result of bacterial inoculation only, and directly precedes the previously described stage I of nodule initiation (Xiao *et al*., 2014), we refer to this stage as stage 0.

**Figure 1.**
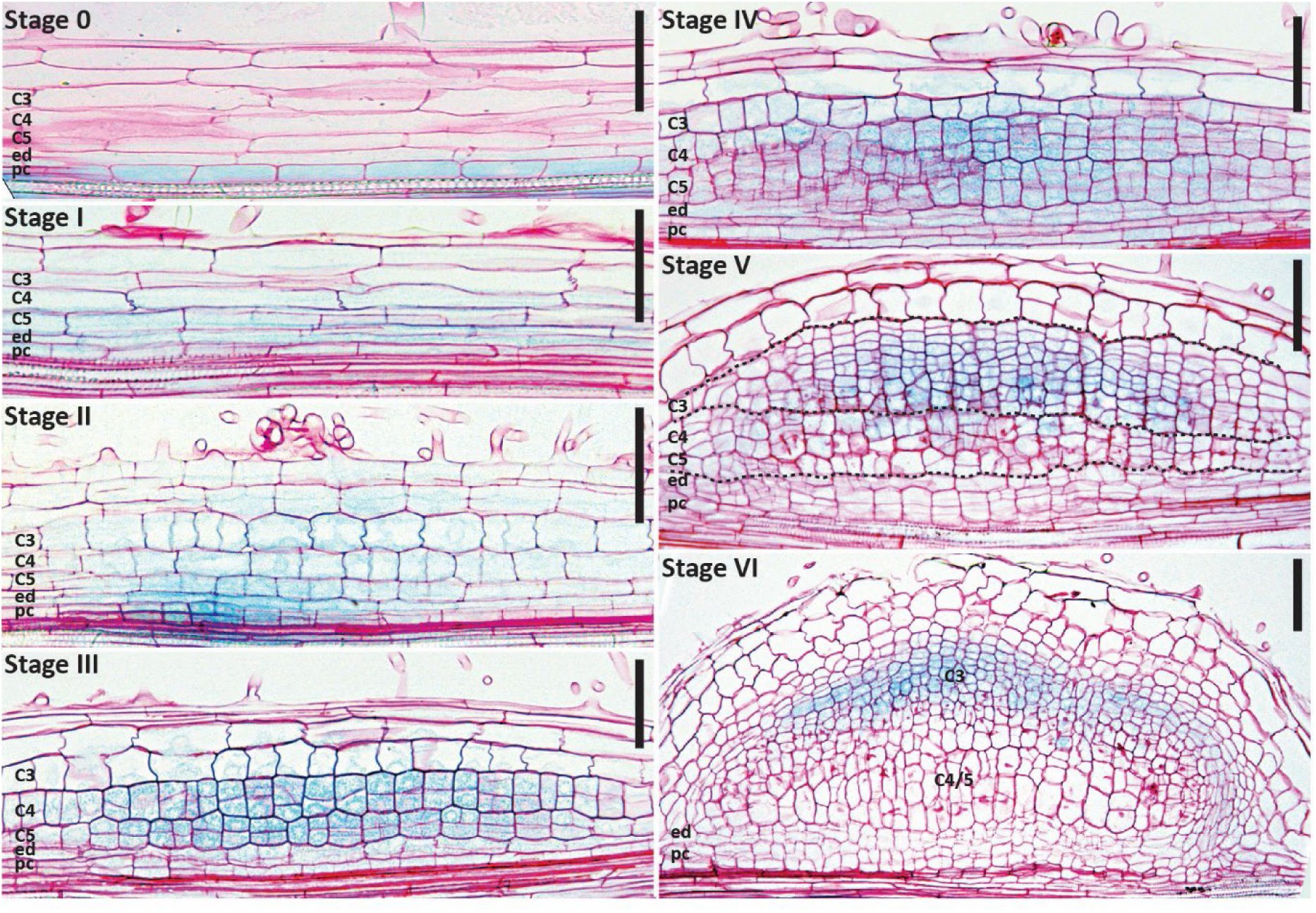
*DR5::GUS* expression dynamics during the different stages of nodule primordium development in Medicago (cv R108). **Stage 0**, *DR5* is activated in pericycle cells preceding their division. **Stage I**, first anticlinal pericycle divisions have occurred, the *DR5::GUS* signal extends to the cortical cell layers prior to their activation. **Stage II**, anticlinal divisions extend to cortex layers 3, 4 and 5, broad DR5 activity is detected in the entire nodule primordia. **Stage III**, periclinal cell divisions in cortex layers 4 and 5, *DR5* is highly expressed in actively dividing cortical cells of these layers. **Stage IV**, periclinal cell divisions in cortex layer 3, DR5 signal extends to the third cortical cell layer. **Stage V**, Cells derived from the C3 layer actively divide to form the future nodule meristem; DR5 activity is highest in these cells. Pericycle, endodermis and cortex layers 4 and 5 have stopped dividing and DR5 activity is no longer detected in the center of these cell layers. **Stage VI**, vasculature bundles are formed, nodule meristem becomes active, DR5 activity is restricted to the nodule vasculature, nodule meristem and infected cells directly adjacent to it. Ep, epidermis; C1-C5, cortical cell layers; ed, endodermis; pc, pericycle. Scale bars 75 μm.

Slightly later in stage I, *DR5* activity was observed in dividing pericycle cells, and not yet dividing cells of the inner root cortex (C4/5). At stage II, *DR5* activity was maintained in these inner layers, and additionally had progressed to the outer cortex layers (C1-3) and epidermis. At stage III and IV, *DR5* was highly active in C4/5 derived cells compared to pericycle and endodermis cells. The latter correlates with decreased or completion of cell divisions in these cell layers at those stages.

During stage V, *DR5* activity in the C4/5 derived cells in the central region of the nodule primordium markedly decreased and was no longer detectable in these cells during stage VI. In contrast, *DR5* was highly active in C3 derived cells, an activity possibly linked to the establishment of the future nodule meristem from these cells.

In addition, in young nodules the infected cells directly adjacent to the nodule meristem, as well as cells of the vascular bundles, displayed relatively high *DR5* activity (Supplemental Fig. S1). This pattern is reminiscent of the pattern of *DR5* activity in young Medicago A17 nodules (Demina *et al*., 2019). Since the majority of experiments that will be described here, were conducted in Medicago Jemalong A17 (A17), we compared *DR5* expression patterns between R108 and A17 during early stages of nodule development. To this end, we introduced *DR5::GUS* to A17 by *Rhizobium rhizogenes* (formerly *Agrobacterium rhizogenes;* (Young *et al*., 2001) *-* mediated hairy root transformation. The composite plants have been grown in perlite for 7 days and then inoculated with *S. meliloti* 2011. At 7 days post inoculation (7dpi) roots were collected and incubated in GUS substrate buffer. Segments of roots with stained nodule primordia at different stages were cut, fixed, embedded in plastic and sectioned. Comparison of R108 sections and sections representing early stages of A17 nodule formation (Supplemental Fig. S2) showed that the *DR5* patterns in both ecotypes were very similar.

#### Selection of auxin-related genes based on their reported expression dynamics during nodule primordium development

Next, we questioned how these *DR5* expression patterns are established. *In planta*, *DR5* activity is the result of auxin gradients established by a dynamic interplay between auxin synthesis and transport (reviewed by Zhao, 2018; Cao *et al*., 2019). First, we focussed on *MtYUCCA* genes (*MtYUCs*) while mining previously published RNAseq data of 10 mm root segments harvested after 3 hrs of Nod factor application (van Zeijl *et al*., 2015), and a time series of *S. meliloti* 2011 spot-applications (Schiessl *et al*., 2019). This revealed that three *MtYUCs* (i.e. *MtYUC1*, *MtYUC2*, and *MtYUC8*) were induced relatively early during nodule initiation, although with different responses over time (Schiessl *et al*., 2019). Moreover, these three *MtYUCs* were expressed in the root susceptible zone before inoculation (van Zeijl *et al*., 2015). A fourth *MtYUC* was found to be expressed during nodule formation, *MtYUC9,* although its expression was only induced after 96hpi (Schiessl *et al*., 2019).

Apart from local biosynthesis, auxin transport is a main contributor to the establishment of the auxin gradients likely responsible for the observed *DR5* expression pattern. *MtPIN1*, *2*, *3*, *4*, and *10* were expressed in the root susceptible zone with *MtPIN4* and *10* contributing >90% of the total *PIN* mRNA pool (van Zeijl *et al*., 2015). The expression of *MtPIN3* and *MtPIN4* was not affected by *S. meliloti* application, *MtPIN1* was only marginally induced from 16hpi onwards*. MtPIN2* and *10* expression was induced after 24hpi, whereas, *MtPIN6* was induced after 36hpi, but dropped to its default expression after 72hpi (Schiessl *et al*., 2019). Additionally, *MtLAX2* was selected as it was the highest expressed *LAX* gene during initiation of nodule formation in Medicago (Roy *et al*., 2017).

Combined, this suggests that auxin synthesis genes *MtYUC1*, *2*, *8*, *9* and auxin transport genes *MtPIN2*, *4*, *6*, *10*, *MtLAX2* are candidate genes likely to be involved in establishing the observed *DR5::GUS* patterns during nodule primordium formation. Therefore, we chose to further analyse the spatiotemporal expression dynamics of these genes.

#### Expression patterns of MtYUCs during nodule initiation and development

To dissect the role of the selected *MtYUC* genes in establishing the auxin patterning during nodule development as reflected by *DR5* activity, we first determined their expression domain in the root susceptible zone. RNA *in situ* hybridization demonstrated that only *MtYUC1* transcripts were clearly detectable in the vasculature of the root susceptible zone (Fig. 2). All four *MtYUC* genes were expressed in the root elongation zone, and the expression of *MtYUC2* and *8* was extended slightly further, to the root differentiation zone (Supplemental Fig. S3). In addition, *MtYUC8* and *9* transcripts were observed in the lateral root cap and columella cells of root tip (Supplemental Fig. S3).

**Figure 2.**
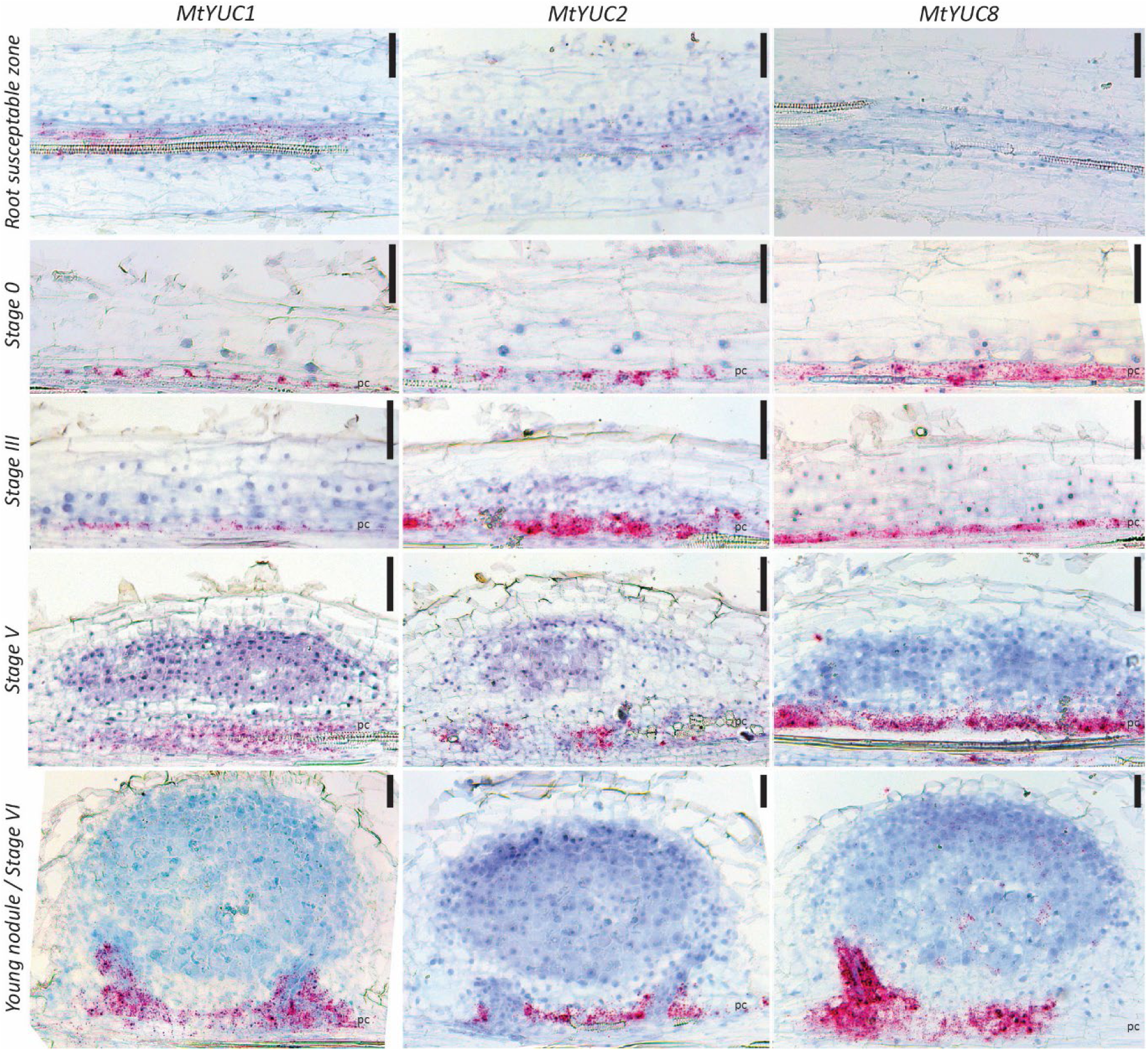
The spatiotemporal expression dynamics of *YUCCAs* in root susceptible zone and during nodule primordium formation. RNA *in situ* hybridizations with *MtYUC1*, *MtYUC2* or *MtYUC8* probe sets on longitudinal sections of root segments and nodule primordia (red dots are hybridization signals, scale bars 75 μm).

To superimpose the spatiotemporal expression dynamics of these *MtYUC*s on the *DR5* pattern, we spot-inoculated Medicago roots and RNA *in situ* hybridization was performed on longitudinal root sections accommodating nodule primordium developmental stages 0 - VI (Fig. 2). For three out of four *MtYUCs*, *MtYUC1*, *2* and *8*, a strong hybridization signal was observed in the pericycle at stage 0. At the stages III to VI, *MtYUC1*, *2* and *8* transcripts remained restricted to the pericycle and the forming vasculature.

This suggests that *MtYUC1*, *2* and *8* could be responsible for the auxin accumulation in the pericycle as visualized by *DR5* activity during nodule initiation. Consequently, it also implies that at later stages the auxin patterns in cortical cells cannot be attributed to the activity of these genes alone. Moreover, the expression of these *MtYUC* genes in the pericycle at later stages does not lead to a strong *DR5::GUS* activity there.

Schiessl and colleagues (2019) reported that the expression of *MtYUC9* was activated at 96hpi. In agreement with this, we did not observe *MtYUC9* transcripts at the earlier stages of nodule primordia development (Supplemental Fig. S4). From stage IV onwards, *MtYUC9* transcripts were detectable at the periphery of the nodule primordium. Probably, these expression domains correlate to the cells of the developing nodule vasculature. At all analysed stages *MtYUC9* expression levels were relatively low (Supplemental Fig. S4). Thereby, the *MtYUC9* expression pattern is also unable to explain the *DR5* activity observed in the root cortex during nodule primordium development from stage II onward.

#### Functional analysis of MtYUCs during nodule initiation

Based on the spatiotemporal dynamics of *MtYUC* expressions during nodule initiation, we hypothesized that local auxin biosynthesis in the pericycle is a requirement for nodule initiation. This raises the question whether inhibition or down-regulation of *MtYUCs* leads to a reduction in nodulation. To test this, we adopted pharmacological and *RNA* interference approaches (Fig. 3).

**Figure 3.**
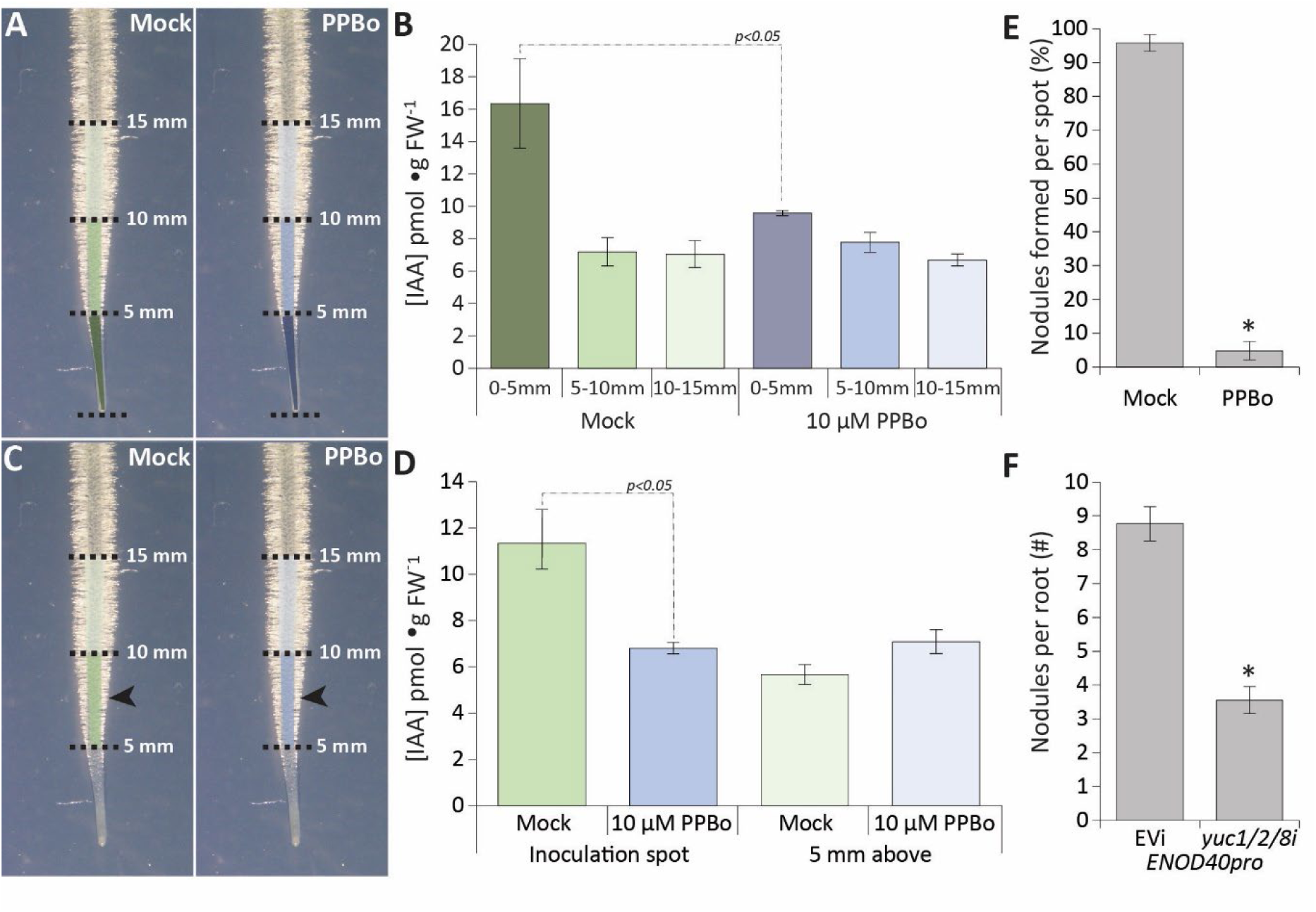
The role of auxin biosynthesis during nodule initiation. **(A, B)** The measurement of auxin (IAA) in 3 zones of roots (root tip, susceptible zone, differentiation zone) after 3 hrs Mock or 10 μM PPBo application. Data are averages of 2 independent experiments (n=8, 2×4), *p*<0.05. **(C, D)** The effect of 24 hrs 10 μM PPBo application on rhizobia induced auxin (IAA) accumulation in the root susceptible zone. Data are averages of 2 independent experiments (n=8, 2×4), *p*<0.05, arrow points at application spot. **(E)** The effect of pharmacological YUCCA inhibition by 10 μM PPBo on the nodulation efficiency of rhizobia spot-applications. Data are averages of 4 independent experiments each containing >10 roots, (n=45), *p*<0.0005. **(F)** Number of nodules formed on *ENOD40*::*YUC1/2/8i (yuc1/2/8i)* transgenic roots compared to EVi transgenic roots. Data are averages of 3 independent experiments each containing >10 roots, (n=50), *p*<0.005. (Bars represent averages, ± standard error, asterisk or p-values demonstrated significant differences between samples).

It has previously been reported that 10 μM of 4-phenoxyphenylboronic acid (PPBo) is very efficient in blocking of the YUC enzymatic activity (Kakei *et al*., 2015). We tested whether addition of 10 μM PPBo prior to inoculation affects nodule initiation. To validate that 10 μM PPBo is able to block IAA biosynthesis in our experiments, we first set out to test its efficiency on Medicago roots. We sampled the lower 1.5 cm of uninoculated roots into 3×5 mm segments (Fig 3A) and determined the IAA concentration in these segments (Fig. 3B). This revealed that IAA levels were approximately 2 times higher in the tip of the root compared to the other two segments (16.34, 7.18, 7.05 pmol•g^-1^ FW respectively). 3 hrs 10 μM PPBo application reduced the level of IAA in the root tip by ∼40%, while not effecting the level of IAA in the two other segments (9.57, 7.78, 6.68 pmol•g^-1^ FW respectively, Fig. 3B). When considering that under these conditions *MtYUC* expression is mostly restricted to the root tip (Fig. 2, Supplemental Fig. S3), this suggests that 10 μM PPBo was able to block local IAA biosynthesis in the root tip while not affecting the IAA derived from the acropetal auxin transport. Next, we treated roots for 24 hrs with 10 μM PPBo or mock, and we subsequently spot-inoculated these roots with rhizobia in the root susceptible zone. These spotted regions and the 5 mm root segments above were harvested for IAA quantification at 24 hpi (Fig 3C, D). In line with previous report (Schiessl *et al*., 2019), rhizobia spot application led to a ∼60% increase in the level of IAA. This increase was not observed when spot application with rhizobia was performed on PPBo treated roots (Fig 3D).

Next, 3-day-old Medicago seedlings were transferred to plates containing either 10 μM PPBo or mock. After 24h, roots of these seedlings were spot-inoculated with rhizobia. At 7 dpi, nodule numbers were scored. This revealed that, on average, nodules formed per spotted root segment were reduced by ∼90% in plants growing on media containing 10 μM PPBo (Fig. 3E). Combined, this suggest that upon Rhizobium applications *Mt*YUC-mediated IAA biosynthesis was blocked by PPBo. As a consequence, IAA levels were reduced, and nodule initiation hampered.

In addition, we created composite Medicago plants bearing transgenic roots in which *MtYUC*s expression was targeted by *RNAi*. It has been shown that in Arabidopsis, the 11 YUC genes have strong functional redundancies, and only higher-order *yucca* mutants display severe developmental defects (Cheng *et al*., 2006, 2007). For this reason, we targeted all three early induced *MtYUCs* (*MtYUC1*, *2* and *8*) simultaneously using one construct. As the transcriptional activation of these *YUCs* is located in the pericycle, we used the *ENOD40* promoter, which is active in the pericycle prior to nodule initiation (Compaan *et al*., 2001) and further activated by Nod factor within 3 hrs in the pericycle (Xiao *et al*., 2014; van Zeijl *et al*., 2015), to drive this *RNAi* construct and test whether such specific knock-down of *YUCs* in this cell layer at the start of nodulation is sufficient to impact nodulation. This was the case, at 3 weeks post inoculation (3wpi), nodulation on *ENOD40:YUCi* transgenic roots was reduced by ∼40% compared to the empty vector control (*EV)* (Fig. 3F).

Apart from blocking rhizobia-induced IAA accumulation in the susceptible zone, 10 μM PPBo also reduced auxin levels in the Medicago root tip. Therefore, we next questioned; is IAA biosynthesis at the tip of the root contributing to the auxin levels required for nodule initiation in the susceptible zone? For this reason, we resectioned the Medicago root tip (3-5 mm) a few minutes prior to *S. meliloti* spot application. Although a relatively harsh treatment, it did not affect nodule initiation compared to nodulation on the intact roots (Fig. 4). This observation indicates that under normal conditions, auxin produced in the root tip is not required for nodule initiation.

**Figure 4.**
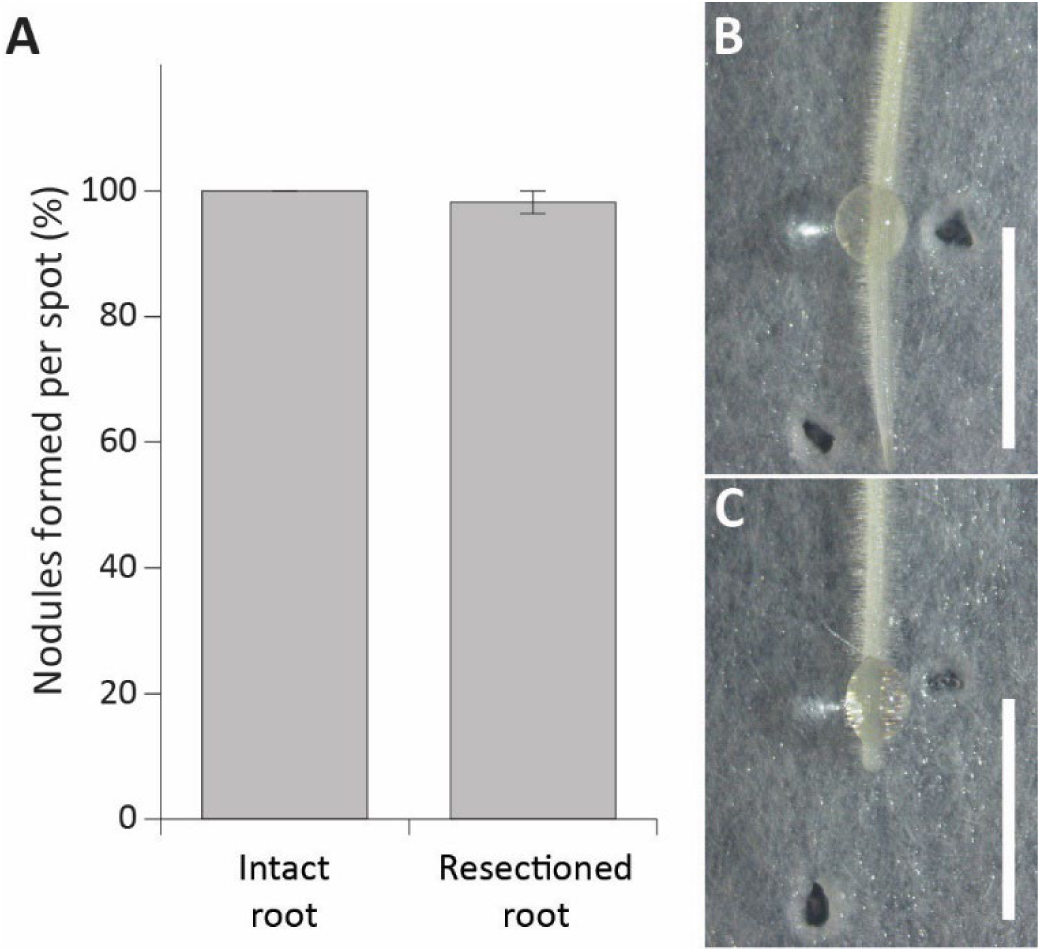
Effect of re-sectioning of the primary root on nodulation. **(A)** Percentage of nodules formed after rhizobia spot-applications on intact or re-sectioned Medicago roots (n>30). (Bars represent averages, ± standard error) **(B)** Application spot of the intact root, **(C)** Application spot and cut site of the re-sectioned root. Scale bar 5mm.

Taken together, *MtYUC1*, *2* and *8* expressions in the pericycle were induced by Rhizobium. All three genes were expressed in pericycle and pericycle-derived cells connected to forming nodule vasculature in the following developmental stages. However, the expression of these *MtYUCs* as well as *MtYUC9* cannot explain the cortical auxin responses as demonstrated by the *DR5* promoter activity in these cells from stage I onwards.

#### Spatiotemporal expression of auxin transporters during nodule primordium development

The observation that the auxin synthesis genes *MtYUC1*, *2*, *8* and *9* are not expressed in cortical tissue during primordium formation raises the question; “how do these cortical cells acquire the auxin as detected by *DR5::GUS*?”. As no other *MtYUCs* were detected during nodule primordium development, an obvious answer to this question is auxin transport. This requires the presence of auxin export carrier *Mt*PIN proteins with such outwards positions of these PINs on the plasma membrane to direct auxin flow from the pericycle towards the cortex. To investigate this, we first determined the spatial expression pattern of the early induced *MtPIN2* and *MtPIN10*. As *MtPIN4* is orthologous to *AtPIN1* (reviewed by Kohlen *et al*., 2018), the main contributor to acropetal auxin transport in the stele of Arabidopsis roots and one highly expressed in the Medicago susceptible zone (van Zeijl *et al*., 2015), we included this gene in our experiments (Fig. 5, Supplemental Fig. S5).

**Figure 5.**
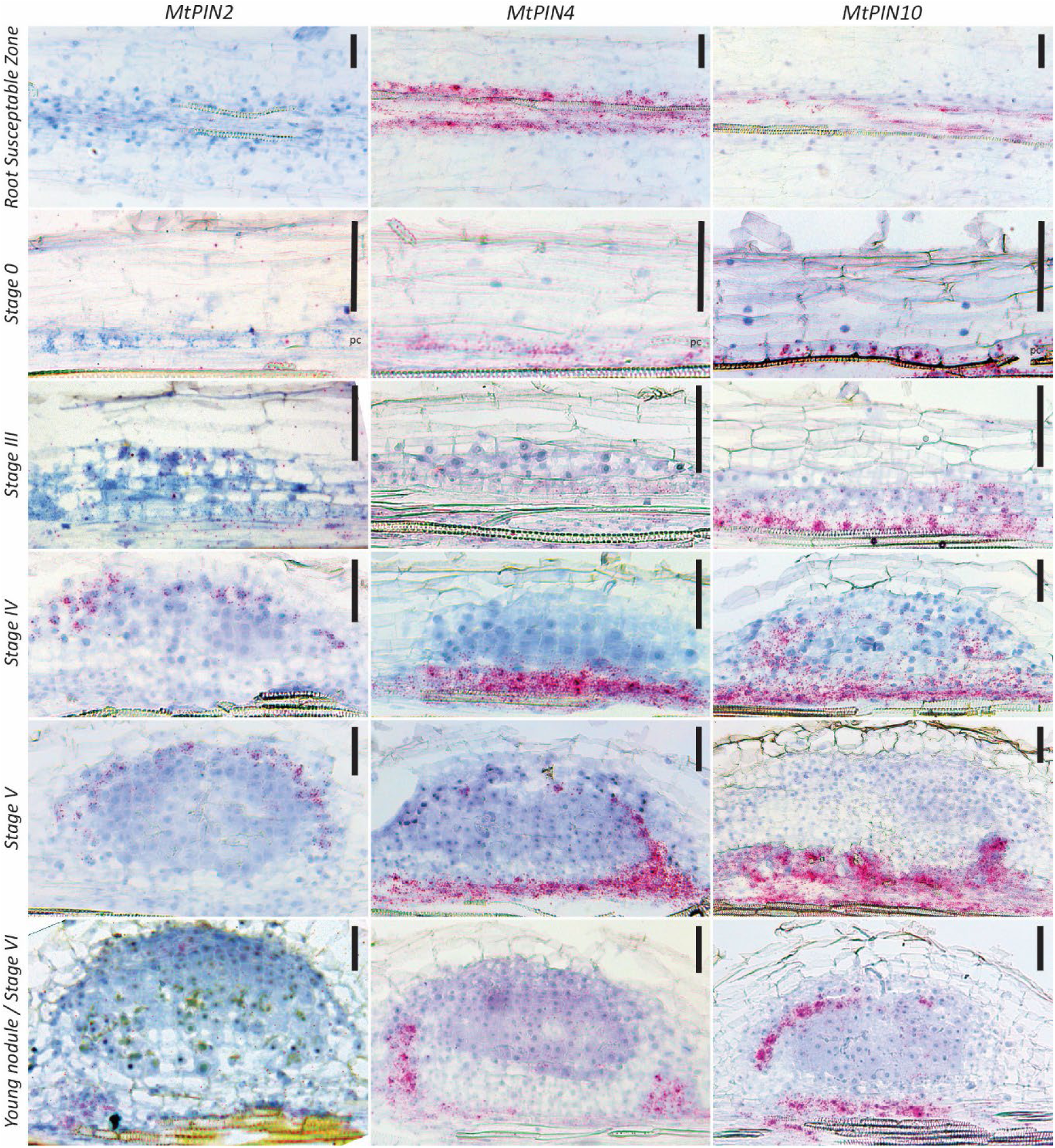
The spatiotemporal expression dynamics of *MtPINs* in the root susceptible zone and during nodule primordium formation. RNA *in situ* hybridizations with *MtPIN2*, *MtPIN4* or *MtPIN10* probe sets on longitudinal sections of root segments and nodule primordia (red dots are hybridization signals, scale bars 75 μm).

In the root susceptible zone, hybridization signals were only observed for *MtPIN4* and *MtPIN10,* in the stele (Fig.5), which agrees with the reported expression levels of these genes (van Zeijl *et al*., 2015). *MtPIN2* transcripts were not detected in susceptible zone. To exclude inefficient *MtPIN2* probe set affinity, we used for RNA *in situ* hybridization longitudinal sections of root tips as a control. In the root tip *MtPIN2, 4* and *10* displayed specific spatial expression patterns (Supplemental Fig. S5), to some extent matching the expression domains of their Arabidopsis orthologous. This demonstrated that *MtPIN2* probe set is capable of detecting *MtPIN2* transcripts, and that this gene is likely not expressed in the root susceptible zone.

Next, we studied the spatiotemporal expression pattern of *MtPIN2*, *MtPIN4* and *MtPIN10* at different developmental stages of nodule primordium development by RNA *in situ* hybridization (Fig. 5). At stages 0-I, *MtPIN2* was not visible in the pericycle. *MtPIN4* was detected in the vasculature and pericycle, and this expression pattern was not different from that in the susceptible zone of un-inoculated plants (Fig. 5). *MtPIN10* transcripts were observed in root vasculature and with highest intensity in the pericycle, suggesting that *MtPIN10* expression was induced in the pericycle at the start of nodule primordium initiation.

At stages III-IV, *MtPIN2* transcripts were seen in both dividing and non-dividing cortical cells, and the developing nodule vasculature. At stage V-VI, *MtPIN2* is expressed mainly in the nodule vasculature and future nodule meristem. Different from *MtPIN2*, *MtPIN4* remained restricted to the pericycle at stage I-III and the forming vasculature at stages IV-VI. *MtPIN10* transcripts were observed also in cortex derived cells at stage III and at stage IV at the periphery of the nodule primordium. At later stages V and VI, *MtPIN4* and *MtPIN10* hybridization signals were visualized mainly in the developing vasculature. Like *MtPIN2, MtPIN10* transcripts were also detectable in the future nodule meristem at stage VI.

*MtPIN6 is* orthologous to the Arabidopsis *PIN6* and has a unique position within the family of PIN auxin transporters as it displays a dual localization, at the plasma membrane and endoplasmic reticulum (Simon *et al*., 2016). As a result, *At*PIN6 is believed to mediate both, auxin polar transport and intracellular auxin homeostasis. Additionally, *At*PIN6 has been demonstrated to be involved in organogenesis (Cazzonelli *et al*., 2013; Simon *et al*., 2016). By using RNA *in situ* hybridization, we could not detect *MtPIN6* in root tip and susceptible zone and at early nodule primordium stages 0-I. However, *MtPIN6* transcripts were visualized in root transition zone and at later time points of nodule development, when nodule vasculature was initiated (Supplemental Fig. S6).

Apart from auxin efflux, auxin influx is a contributing factor to establish directional auxin transport (Bennett *et al*., 1996). The auxin import carrier *MtLAX2* is highly induced during the early stages of nodule development (de Billy *et al*., 2001; Roy *et al*., 2017; Schiessl *et al*., 2019). We performed *in situ* RNA hybridisation on the Medicago root, including the root tip and susceptible zone. Our results confirmed previously published expression pattern of *MtLAX2* that was detected in the lateral root cap, in some of columella cells and developing vasculature (de Billy *et al*., 2001) (Supplemental Fig. S7). In addition, strong *MtLAX2* hybridization signals were detected in the vasculature of the susceptible zone. To resolve *MtLAX2* expression dynamics during nodule primordium formation, we performed RNA *in situ* hybridization on longitudinal sections of spot-inoculated root segments. This revealed that, at stage 0, *MtLAX2* transcripts were located in the root vasculature and pericycle (Supplemental Fig. S8). At stage III, *MtLAX2* expression was detected mainly in the pericycle, forming nodule vasculature and, at later stages, in a future nodule meristem albeit at very low levels.

Taken together, our spatiotemporal studies on auxin transporter gene expression revealed that *MtLAX2*, *MtPIN4* and *MtPIN10* could be involved in local auxin accumulation at stage 0 and I in the pericycle, while *MtPIN2*, *MtPIN6* and *MtPIN10* could participate in creating the auxin pattern in cortical cells.

#### Mt*PIN10* dynamics during nodule formation

If auxin biosynthesis at stage I and II of nodule primordium initiation does not occur locally in the cortex and both *MtPIN4* and *MtPIN10* are expressed in the pericycle, it implies that auxin might be produced in the pericycle and actively transported towards the cortex. To determine if cellular localization of the expressed *Mt*PIN proteins could facilitate such transport, we created a *MtPIN10::PIN10-eGFP* (*Mt*PIN10-GFP) protein fusion. We attempted to create a similar construct for *Mt*PIN4, but unfortunately without success.

First, we tested whether this construct would mimic the expected expression domains. Indeed, composite plants baring *Mt*PIN10-GFP transgenic roots showed a similar expression pattern in the root tip (Supplemental Fig. S9) as observed by RNA *in situ* hybridisation for *MtPIN10* (Supplemental Fig. S5). However, no GFP signal was detected in the susceptible zone for *Mt*PIN10 where this gene is expressed (Supplemental Fig. S9).

Next, we spot-inoculated roots transiently expressing *Mt*PIN10-GFP with *S. meliloti 2011*. Root segments were harvested at different time points after inoculation, longitudinal hand-made sections through the vasculature were made, and immediately observed under confocal microscope. To determine subcellular *Mt*PIN10 localization, we analysed ∼5-7 primordia for each stage and representative images are shown (Fig. 6).

**Figure 6.**
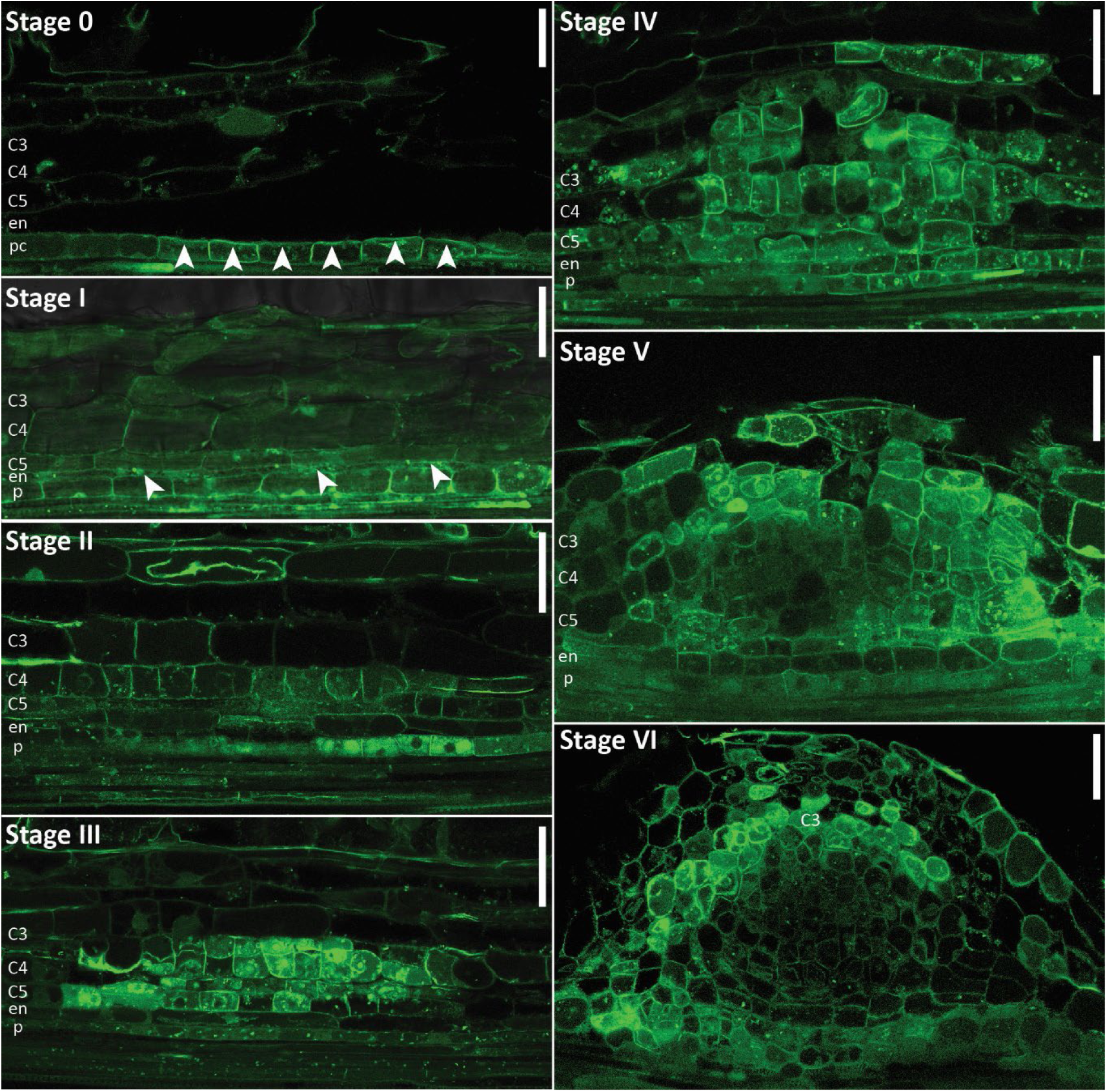
*Mt*PIN10 localization during Medicago nodule primordium formation. **Stage 0,** *Mt*PIN10-GFP is detected in pericycle cells, partially orientated towards the cortex (arrowheads). **Stage I,** *Mt*PIN10-GFP is extended to the endodermis (arrowheads). **Stage II,** *Mt*PIN10 levels are reduced in the pericycle. *Mt*PIN10 levels are increased in cortical cells with no apparent polarity. **Stage III,** *Mt*PIN10 is present in pericycle only at low levels. High *Mt*PIN10 levels are detected in dividing cortical cells with no clear polarity. MtPIN10 is also detected in the root vasculature positioned on the plasma membranes towards the root tip. **Stage IV,** *Mt*PIN10 is positioned towards the centre of the primordia in the cells located at the primordium periphery and no clear polarity is visible in the cells located at the centre of the primordium. **Stage V/ VI,** *Mt*PIN10 is hardly detectable in the central part of the primordia, at the periphery, in developing nodule vasculature cells, it is positioned towards the (future) nodule meristem (scale bars, 75 μm).

At stage I, *Mt*PIN10 was clearly visible in the pericycle at the site of nodule primordium induction, and, although not exclusively, in part localized towards the endodermis and root cortex (arrowheads). At stage II, *Mt*PIN10 was visible in the endodermis and inner cortex. At stage III *Mt*PIN10 was localized in the dividing cortical cells, and later at the primordium periphery positioned towards the centre of the activated cortical cells. At later stages (V and VI) *Mt*PIN10 localized in the periphery of the nodule primordia, likely the nodule vasculature. Here, subcellular localization of *Mt*PIN10 was orientated towards the nodule meristem. Currently, we do not know whether *Mt*PIN4 follows a similar localization pattern compared to *Mt*PIN10.

#### Functional analysis of MtPINs during nodule initiation

A reported Medicago *pin2* mutant did not show a nodulation-related phenotype (Ng *et al*., 2020). These observations imply that there is a certain functional redundancy of *MtPINs*. Such redundancy is known in Arabidopsis roots where PIN proteins exhibit synergistic interactions mechanism when the loss of a specific PIN protein is compensated by auxin dependent ectopic expression of its homologues (Vieten et al., 2005). Two other mutant lines have been isolated and described in Medicago, loss-of-function *pin10/slm1* (SMOOTH LEAF MARGIN1) (Peng and Chen, 2011; Zhou *et al*., 2011*a*,*b*). These mutant lines display severe pleiotropic phenotypes in different organs including leaf and flower development. In addition, both lines are sterile. For this reason, these mutants must be maintained as heterozygous and for neither mutant a nodule phenotype has been described.

We inoculated Medicago *pin10-1* mutant plants with rhizobia and analysed nodules at 3wpi. The *pin10-1* mutation had no significant effect on the number of nodules formed per plant compared to wild-type R108 (Fig. 7A). This observation suggests that *MtPIN10* either plays no role during nodulation, or its involvement is masked by a functional redundancy. Indeed, *MtPIN4* has a similar expression pattern as *MtPIN10* (Fig. 5), and therefore might function redundantly to *MtPIN10*. To test this, we generated *RNAi* construct to knock-down *MtPIN4* and *MtPIN10* simultaneously.

**Figure 7.**
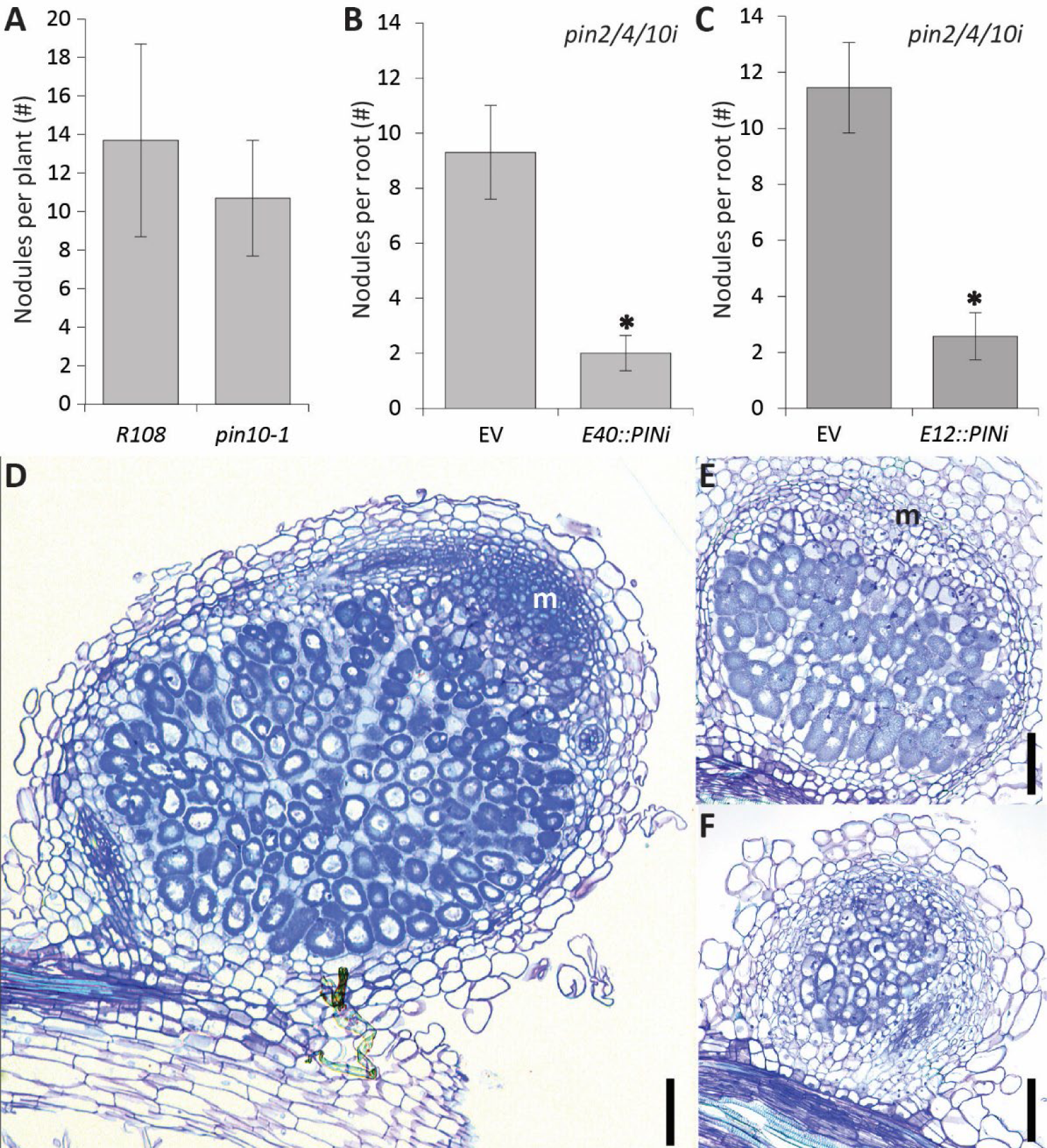
The role of auxin transport during nodulation in Medicago. (A) The number of nodules formed on roots of *pin10-1* compared to wild type (R108), n=40. **(B)** The number of nodules formed on roots expressing an Empty Vector control (EV), or *ENOD40*::*PIN2/4/10i* (*E40::PINi*) construct, n=30. **(C)** The number of nodules formed on roots expressing an Empty Vector control (EV), or *ENOD12*::*PIN2/4/10i* (*E12::PINi*) construct, n=30. Bars represent averages, ± standard error, asterisk demonstrated significant differences between samples, *p*<0.05). **(D-F)** Representative pictures of **(D)** an *EV* control nodule containing a fully developed nodule meristem, **(E)** *an E12:PINi* nodule with a short and underdeveloped meristem, and **(F)** an *E12:PINi* nodule without a meristem; scale bars 100µm, m=meristem.

We used the *ENOD40* promoter to silence these genes at the very beginning of nodule initiation (Compaan *et al*., 2001). The introduction of this construct *in planta* by hairy root transformation reduced nodule number by ∼70% compared to the *EV* control (Fig. 7B). This demonstrates that expression of *MtPIN4* and *MtPIN10* in the pericycle is required for nodule initiation. However, as the *ENOD40* promoter is also active beyond stage VI of nodule organogenesis (Crespi *et al*., 1994), our results cannot exclude PINs to be important during later stages (Fig. 5). Therefore, we silenced *MtPIN4*, *MtPIN10* and *MtPIN2* (also expressed at later stages of nodule primordia formation) under a the symbiosis specific promoter *ENOD12* (Li *et al*., 2021). Introduction of this construct led to a similar reduction in nodule numbers compared to the *ENOD40* promoter (Fig. 7B-C). Moreover, the few nodules formed were small compared to wild type, and plastic sections revealed that they rarely (7 out of 22 vs 22 out of 25 for the EV control) formed a meristem (Fig. 7D-F, table 1). Combined, this shows that downregulation of PINs under control of the ENOD40 and ENOD12 promoter have a similar effect on nodule numbers. This suggests that downregulation of these *PINs* prior to rhizobium application might not have an effect on nodule numbers.

**Table 1:** *E12::PINi nodule ontology*

| Line | Nodules sectioned | Normal | Small or no meristem |
| --- | --- | --- | --- |
| EV | 25 | 22 | 3 |
| *E12::PINi* | 22 | 7 | 15 |

## DISCUSSION

Here, we presented dynamics of genes and proteins involved in auxin biosynthesis and transport and propose their role in establishing the *DR5* expression patterning during sequential stages of nodule formation. From this, the following picture emerges with respect to how the auxin landscape links to nodulation; *DR5* activity is concentrated in pericycle cells preceding their first division, and in a wave like motion, only a few cells wide, the *DR5* activity moves through the endodermal and cortical cells that give rise to the nodule primordium, vasculature and ultimately the nodule meristem. Based on this *DR5* patterning, four major events during nodule development in which auxin likely plays an important role can be defined as: pericycle activation, cortex activation, vascularisation, and meristem establishment.

### Local auxin accumulation in the pericycle is a prerequisite for nodule initiation

The models presented by Deinum and colleagues (2012, 2016) gave priority to a local reduction of auxin efflux (PINs) to explain the auxin maximum preceding nodule formation (Xiao, 2015). It has been suggested that this reduction in transport capacity leads to a local increase of auxin levels in the pericycle and inner cortex at the site of nodule primordium initiation (Mathesius *et al*., 1998*b*,*a*; Ng and Mathesius, 2018). However, a direct comparison between auxin transport inhibition and *S. meliloti* application is difficult as the auxin response patterns and timing of the initiation between these two treatments are different (Ng and Mathesius, 2018).

We demonstrate that a local auxin transport network of *MtPIN4/10* and *MtLAX2* are expressed in the stele of the root susceptible zone, and that *MtLAX2* is highly expressed in the pericycle at the start of nodule initiation. It is possible that together these auxin transporters create a conduit for an auxin flow from the stele into the pericycle. However, our data show that during the first stages of nodule primordium development (stage 0-I) the expression of the auxin biosynthesis genes *MtYUC1*, *2* and *8* correlates with the mitotic re-activation of the few cells in the pericycle that initiate a nodule primordium. Pericycle specific *RNAi* against these *MtYUCs (ENOD40::YUCi)* and pharmacological applications with the YUCCA inhibitor ppBO demonstrates that local activity of *MtYUC1*, *2 and 8*, and therefore likely local auxin biosynthesis, in the pericycle is crucial for nodule initiation and further progression. Additionally, we have shown that re-sectioning of the root tip, and thus removal of the auxin located there, had no effect on nodule initiation. This demonstrates that basipetal auxin transport, likely facilitated by *Mt*PIN2, is not involved in nodule initiation. This result is in line with published data on *Mtpin2* mutant that does not display any nodule phenotype (Ng *et al*., 2020).

Our findings that the addition of PPBo, while blocking local auxin biosynthesis and nodule initiation, has no effect on the auxin pool originating from the shoot suggests that acropetal auxin transport might be less important in delivering the auxin required for the establishment of the nodule-initiating auxin maximum. Nevertheless, we cannot exclude that auxin transport from the stele to the pericycle is needed to induce *Mt*YUCs mediated local auxin biosynthesis in a feed-forward fashion. Indeed, it has been demonstrated that in Arabidopsis, local auxin biosynthesis and long-distance auxin transport are collaborating in regulation of the auxin homeostasis required for main and lateral root growth (Marchant *et al*., 2002; Laskowski *et al*., 2008; Chen *et al*., 2014; Guo *et al*., 2014; Tang *et al*., 2017). Thus, our data suggest that a similar scenario could be underlying nodule initiation. On the other hand, it is possible that auxin transport attenuation is only needed to sustain an auxin maximum created by local auxin biosynthesis in the pericycle.

### Is pericycle derived auxin a trigger for mitotic reactivation of cortical cells?

As the expression of *MtYUC1*, *2*, and *8* during the early stages of nodule initiation is restricted to the pericycle, their spatial expression cannot explain the *DR5* pattern beyond stage I. The only other *MtYUC* that is reported to be expressed during nodule primordia formation is *MtYUC9*. However, this gene is only expressed at later stages of nodule initiation (Schiessl *et al*., 2019). Our *in situ* hybridisation experiments demonstrated that *MtYUC9* expression is only detectable from stage IV onward, and only at the periphery of the nodule primordium. Therefore, it is unlikely that the expression of *MtYUC9* is responsible for the auxin required to initiate the cortical *DR5* activity during the preceding stages II and III.

If local auxin biosynthesis does not occur in the cortex, it cannot be directly linked to the *DR5* activity preceding early cortical cell divisions. A plausible explanation could be that the observed *DR5* expression pattern is the result of auxin transported into the cortex, likely from the pericycle. If so, this suggests that the activated pericycle cells are serving as an auxin source. Transport towards the cortex could be mediated by auxin efflux- and influx carriers. *MtPIN4*, *MtPIN10* and *MtLAX2* are expressed in the pericycle during the early stages of nodule initiation, and at least for *Mt*PIN10 protein, we have shown a localization in the pericycle and endodermis that could create an auxin transport channel from the pericycle through the endodermis into the cortex. Our spatiotemporal studies on auxin transporter gene expression reveals that, although *MtPIN2* and *MtPIN6* are expressed in the Medicago root, and their level of expression is regulated during nodule initiation, they are expressed at different positions or later stages of nodule primordia development. Combined with results of Roy et al., (2017), we propose that *Mt*PIN4 and 10, in concert with *Mt*LAX2, are likely candidates to facilitate this putative auxin flux from the pericycle into the cortex.

### Is the nodule meristem an auxin sink or source?

At stages V and VI, the expression of *MtYUC1, 2* and *8* is confined to the base of the nodule and nodule vasculature, while *DR5* activity is highly concentrated in the C3 derived cortical cells where the nodule meristem is about to be formed. As such, also at this stage *DR5* activity and local auxin biosynthesis does not fully match.

In Arabidopsis, locally produced auxin by *At*YUCs is a principal factor for vascular strand formation (Cheng *et al*., 2006; Cao *et al*., 2019). The expression of *MtYUC1*, *2* and *8* in the developing vasculature combined with a weak *DR5* activity indicates that in Medicago *Mt*YUCs might fulfil a similar role in vascular strand formation. However, the strong *MtYUC1*, *2* and *8* expression at the base of the nodule, in a cell layer likely derived from pericycle, does not lead to *DR5* activity in these cells. This could mean two things. Either, as the synthetic auxin reporter *DR5* is based on a single auxin response factor binding motif (i.e. motif bound by AtARF1) (Ulmasov *et al*., 1997), the observed *DR5* activity might not reflect the total auxin dynamics. On the other hand, as *DR5* activity requires auxin to be nuclear localized (reviewed by Yu *et al*., 2022), the absence of a strong *DR5* signal here could therefore imply that most of the auxin produced in the cells at the base of the nodule is immediately transported away. This putative auxin flow towards the forming nodule meristem would explain the *DR5* activity observed in the C3 derived cells. It is possible that auxin, synthesized in the base of the nodule, the nodule vascular bundles, or even from the acropetal auxin transport stream, is translocated through the nodule vasculature towards the future nodule meristem.

Such a flux of auxin into the nodule meristem would imply that the developing vasculature is not the endpoint of the biosynthesised auxin, but a passage. The auxin flowing through could be the signal for connecting the future meristem to the central vasculature.

Indeed, *MtPIN2* and *10* genes are expressed in the nodule vasculature as well as in the C3 derived cells forming the future nodule meristem, and therefore likely candidates to function in such auxin transport. Medicago mutants that fail to develop proper nodule vascular bundles as *Mtnoot1/2* (Magne *et al*., 2018; Shen *et al*., 2020) or *Mtlin4* (Guan *et al*., 2013), while being impaired in nodule meristem development or maintenance are supporting the hypothesis that nodule vascular bundles are involved in nodule meristem development.

If so, this would suggest that the auxin content of the nodule meristem is regulated through transport of auxin produced elsewhere. We demonstrated that, when the level of *MtPIN2*, *4* and *10* is reduced during nodule initiation, possibly in the forming nodule vasculature, the formation of the nodule meristem is hampered. Combined, this suggests that auxin transport is needed for the nodule meristem to be formed, and as a consequence the nodule meristem is likely an auxin sink. This auxin can either be transported directly through the C5-C4 layers, or through the developing nodule vasculature.

Based on the canalisation hypothesis the latter is the most likely option. This hypothesis states that a high source of auxin finds its way to an auxin sink, and in doing so paves the way for the formation of a vascular strand (reviewed by Hajný *et al*., 2022). During leaf vascular patterning, the tip of the leaf primordia is an auxin source, and as a consequence vascular strand development progresses from the tip of the leaf towards the base (Scarpella *et al*., 2006; reviewed by Biedroń and Banasiak, 2018). If the nodule meristem is indeed an auxin sink, this canalisation hypothesis implies that vascularisation would progress from the base of the nodule towards the nodule meristem, which would be in contrast to leaf vascularization. Such a nodule meristem can be considered a non-autonomous acting meristem. This would be an elegant way for the plant to regulate meristem functioning and as a direct result, nodule lifetime. Also, this might indicate that in determinate nodules, like in Lotus and soybean, meristem formation is terminated early onwards because of lack of auxin transported into the meristem.

It has been proposed that root nodules are evolutionary derived from lateral roots (reviewed by Sprent, 1989; and Hirsch *et al*., 1997; Franssen *et al*., 2015; Schiessl *et al*., 2019; Soyano *et al*., 2019). During lateral root initiation, the lateral root meristem starts its life as an auxin sink, as shoot-derived auxin is needed for lateral root initiation (Bhalerao *et al*., 2002; Marchant *et al*., 2002). Between stage III and V of lateral root initiation (reviewed by Du and Scheres, 2018), the lateral root primordia becomes autonomous from the shoot derived auxin as the lateral root meristem starts producing auxin itself (Ljung *et al*., 2005). Our results indicate that this switch from sink to source does not occur in the nodule meristem. This inability of the nodule meristem to become an auxin source should therefore be considered an additional key difference between these two developmental processes.

In conclusion, our data suggest that a precise spatiotemporal regulation of auxin outputs are guiding multiple stages of nodule primordium initiation and development. This spatiotemporal regulation is likely the result of a balanced interplay between local auxin biosynthesis and transport. We propose that during the early stages of nodule initiation, auxin is probably biosynthesized in the pericycle, from where it is transported into the cortex to re-activate cell divisions in the C4 and C5 layers. At later stages, when the auxin flow from the pericycle into cells derived from these C4 and C5 layers diminishes, these cells stop dividing and start to differentiate. At the same time, auxin biosynthesized elsewhere (possibly in the pericycle, developed nodule vasculature and/or acropetal auxin transport stream in the main root) is needed to initiate meristem formation and later ensure its persistence. At this moment, the nodule meristem is a sink, draining the surrounding cells of their auxin. Proximal to the nodule meristem, in the cells derived from the C4 layer and later from the nodule meristem, this auxin reduction triggers cell differentiation leading to the infection zone.

## METHODS

### Plant material and bacterial strains

Medicago R108 seedlings were used to make the stable *DR5::GUS* transgenic line by using *Rhizobium radiobacter* (formerly *Agrobacterium tumefaciens*) (strain AGL1) according to the protocol described by (Chabaud *et al*., 2003). Medicago Jemalong A17 plants were used to generate *A. rhizogenes* (strain MSU440) mediated transgenic roots as previously described by Limpens et al. (2004) for RNAi and protein GFP fusion constructs. *Mtpin10-1/slm1 (*NF3969) were previously described by (Zhou *et al*., 2011*a*). Surface-sterilization and germination of Medicago seeds were performed as previously described by Limpens et al. (2004). *Sinorhizobium meliloti 2011* was used to induce root nodule formation on seedlings growing in perlite or on plates with low nitrate Fähraeus (Fä) medium (Fähraeus, 1957).

### Constructs

DNA fragments were amplified from Medicago genomic DNA using primer combinations listed in Table S1 and Phusion*™* High*-*Fidelity DNA Polymerase (Finnzymes). To create the constructs, pENTR™/D-TOPO® Cloning Kits (Invitrogen) and Gateway® technology (Invitrogen) were used to generate the entry clones for RNAi constructs, promoter-GUS and PIN10-GFP constructs. First, 14 synthetic DR5 DNA fragments (Ulmasov *et al*., 1997) were introduced in the entry clone. Then, the entry vector was recombined into Gateway®-compatible binary vector pKGW-RR, that contains the GUS reporter gene and *AtUBQ10::DsRED1* as a selection marker (Limpens *et al*., 2004), by using Gateway® LR Clonase® II enzyme mix (Invitrogen). PCR fragments of about 400-600 bp of single *MtYUCs* and *MtPINs* genes for RNAi constructs were generated on cDNA made from Medicago nodule or root RNA. These fragments were combined by subsequential PCR steps using primers with a complementary 15 bp overhang to generate one amplicon of two or three *DNA* fragments. The final DNA fragments were cloned into pENTR-D-TOPO and recombined into the Gateway-compatible binary vector pK7GWIWG2(II)-UBQ10::DsRED driven by *ENOD40* promoter (Compaan *et al*., 2001; Limpens *et al*., 2004) to create the final RNAi construct. For *Mt*PIN10GFP fusion constructs, the promoter and first part of the DNA fragment was introduced into Gateway® donor vector pENTR2-1, eGFP DNA fragment without start codon were introduced into pENTR1-2 and the rest of the gene including the terminator was introduced into pENTR2-3, using Gateway® BP Clonase® II enzyme mix. These entry vectors were recombined into Gateway®-compatible binary vector pKGW-RR-MGW containing *AtUBQ10::DsRED1* as a selection marker using Gateway® LR Clonase® II Plus enzyme mix (Invitrogen). Insertion sites for the eGFP reporter in *PIN* genes were selected based on functional AtPINs::GFP protein fusion constructs in Arabidopsis (Xu and Scheres, 2005), which are at position 1237 and 1565 for *MtPIN2* and *MtPIN10*, respectively. All used primers are listed in Table S1.

### Tissue embedding, sectioning and staining

Root segments were fixed in 4% paraformaldehyde (w/v), 5% glutaraldehyde (v/v), 0.05 M sodium phosphate buffer (pH7.2) at 4 °C overnight. The fixed material was dehydrated in an ethanol series and subsequently embedded in Technovit 7100 (Heraeus Kulzer) according to the manufacturer’s protocol. Sections (7 µm) were made by using a RJ2035 microtome (Leica Microsystems, Rijswijk, The Netherland), stained 5 min in 0.1% Ruthenium Red (Sigma, Germany). Sections were analysed by using a DM5500B microscope equipped with a DFC425C camera (Leica Microsystems, Wetzlar, Germany).

### Suppression of YUCCAs expression by 10μM ppBO

The germinated Medicago seeds were placed on Fa plates for 3 days and then seedlings were transferred to new plates containing Fa medium with either 10 μM PPBo or mock (Kakei *et al*., 2015). After 24 hrs roots of these seedlings were spot-inoculated with rhizobia (OD_600_=0.02). Plants were grown further for 4 more days (finally 6-7 dpi) to form nodules.

### Auxin quantification

Auxin was quantified through Liquid Chromatography-Mass Spectrometry (LC-MS/MS). Medicago roots were collected and flash frozen in liquid nitrogen. Collected tissue was ground to a fine powder at -80°C using 3 mm stainless steel beads at 50 Hz for 2*30 seconds in a TissueLyser LT (Qiagen, Germantown, USA), and between 10-15 mg of ground tissue per sample was used for auxin extraction. Samples were extracted with 1 mL of cold methanol containing [phenyl ^13^C_6_]-IAA (0.1 nmol/mL) as an internal standard in a 2-mL eppendorf tube and purified as previously described (Ruyter-Spira *et al*., 2011). Samples were filtered through a 0.45 μm Minisart SRP4 filter (Sartorius, Goettingen, Germany) and measured on the same day. Auxin was analyzed on a Waters Xevo TQs tandem quadruple mass spectrometer as previously described (Ruyter-Spira *et al*., 2011; Gühl *et al*., 2021).

### RNA in situ hybridization

RNA *in situ* hybridization was conducted using Invitrogen ViewRNA ISH Tissue 1-Plex and 2-Plex Assay kits (ThermoFisher Scientific) using the manufacture’s protocol optimized for Medicago root and nodules sections (Kulikova *et al*., 2018). RNA ISH probe sets were designed and synthesized at ThermoFisher Scientific. Assay ID of used probe sets are presented in Table S2. As a negative control for each hybridization run one slide was used without any probe set. MtNFYA1 probe set Type 6 was used as a marker for nodule primordium initiation (blue hybridization signals). The images were taken with an AU5500B microscope equipped with a DFC425c camera (Leica).

### Detection of MtPIN10GFP

Transgenic roots or root nodules were manually sectioned in a longitudinal direction, mounted on microscope slides and immediately observed under a Leica SP8 confocal microscope. GFP is visualized by excitation at 488 nm and by detection at 505–530 nm.

### Statistics

When appropriate, data were subjected to the Student’s t test (Microsoft Excel). All other data were subjected to one-way ANOVA. Individual differences were then identified using a post hoc Tukey test (P < 0.05). All analyses were performed using SAS_9.20 (http://www.sas.com/).

## AUTHOR CONTRIBUTIONS

Conceptualization, HF, TB, OK, and WK; Methodology OK, and WK; Investigation, TTX, DS, SM, JL, AvS, HF, OK, and WK; Writing – Original Draft, TTX, HF, TB, OK, and WK; Writing – Review & Editing, all authors; Writing – final version, HF, OK, and WK; Funding Acquisition, TB, and WK; Supervision, OK, and WK.

## ACKNOWLEDGEMENTS

This research is funded by European Research Council (ERC-2011-AdG-294790) to TB, NWO-VENI (863.15.010) and NWO-VIDI (VI.Vidi.193.119) to WK, and Graduate School ‘Experimental Plant Science’.

## Supplemental Figures

**Supplemental Figure S1.**
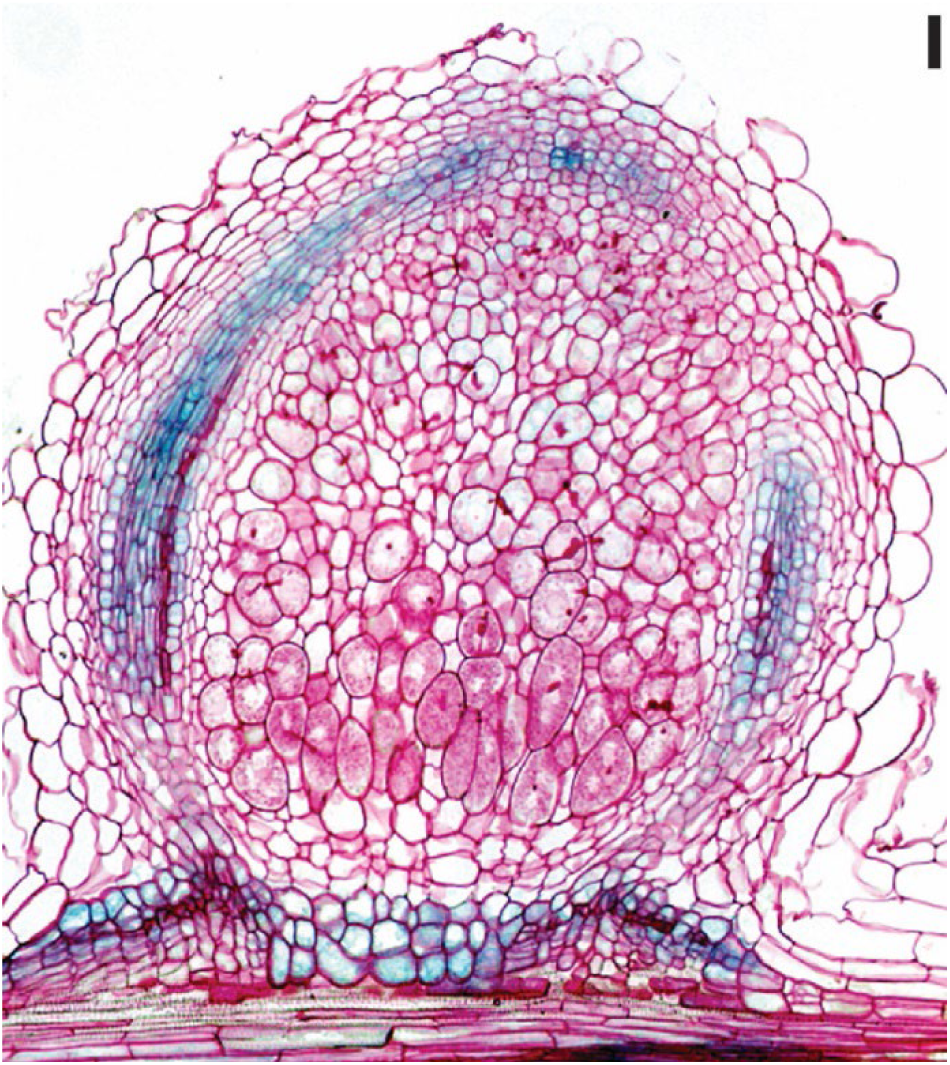
*DR5::GUS* expression pattern in Medicago R108 nodule. (Scale bars 75 μm).

**Supplemental Figure S2.**
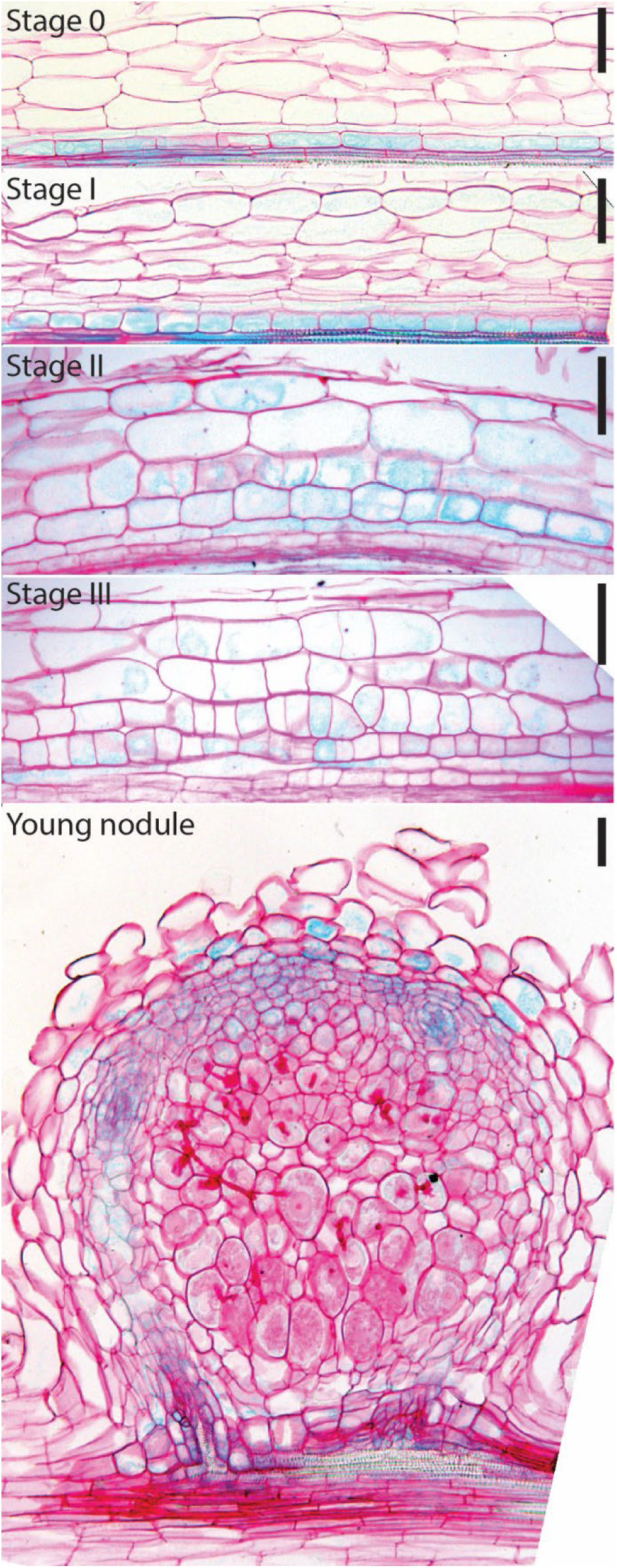
Dynamics of *DR5::GUS* expression patterns during nodule development in Medicago A17. Scale bars 75 μm).

**Supplemental Figure S3:**
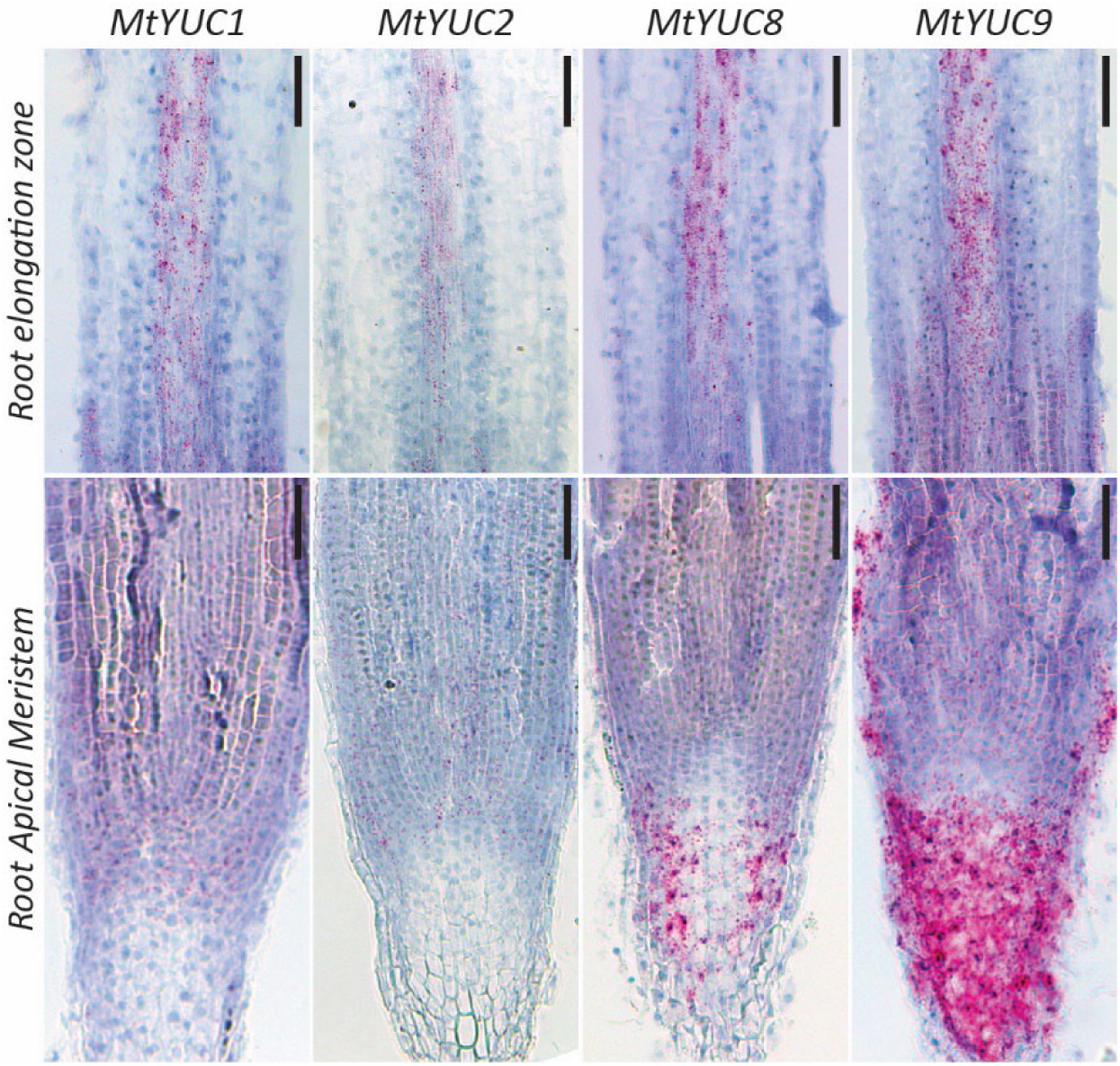
The spatiotemporal expression patterns of *YUCCAs* in root tip of Medicago A17. RNA *in situ* hybridizations with *MtYUC1*, *MtYUC2*, *MtYUC8* or *MtYUC9* probe sets on longitudinal sections of the root tip including elongation zone and root meristem (red dots are hybridization signals, scale bars 75μm).

**Supplemental Figure S4:**
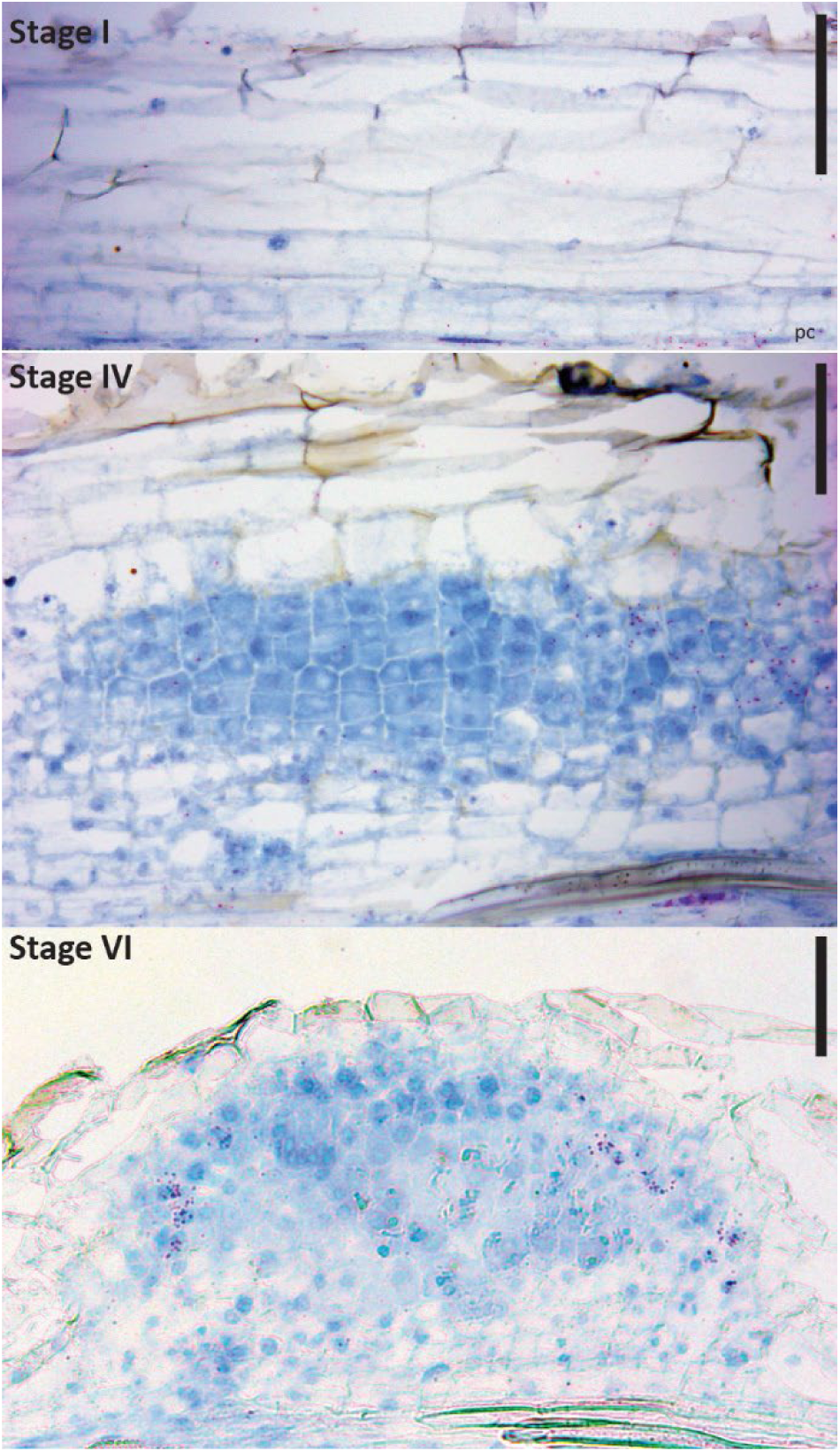
The spatiotemporal expression patterns of *MtYUC9* during nodule primordium formation in Medicago A17. RNA *in situ* hybridization with *MtYUC9* probe set on longitudinal sections of nodule primordia stages I, IV and V (red dots are hybridization signals, scale bars 75 μm).

**Supplemental Figure S5:**
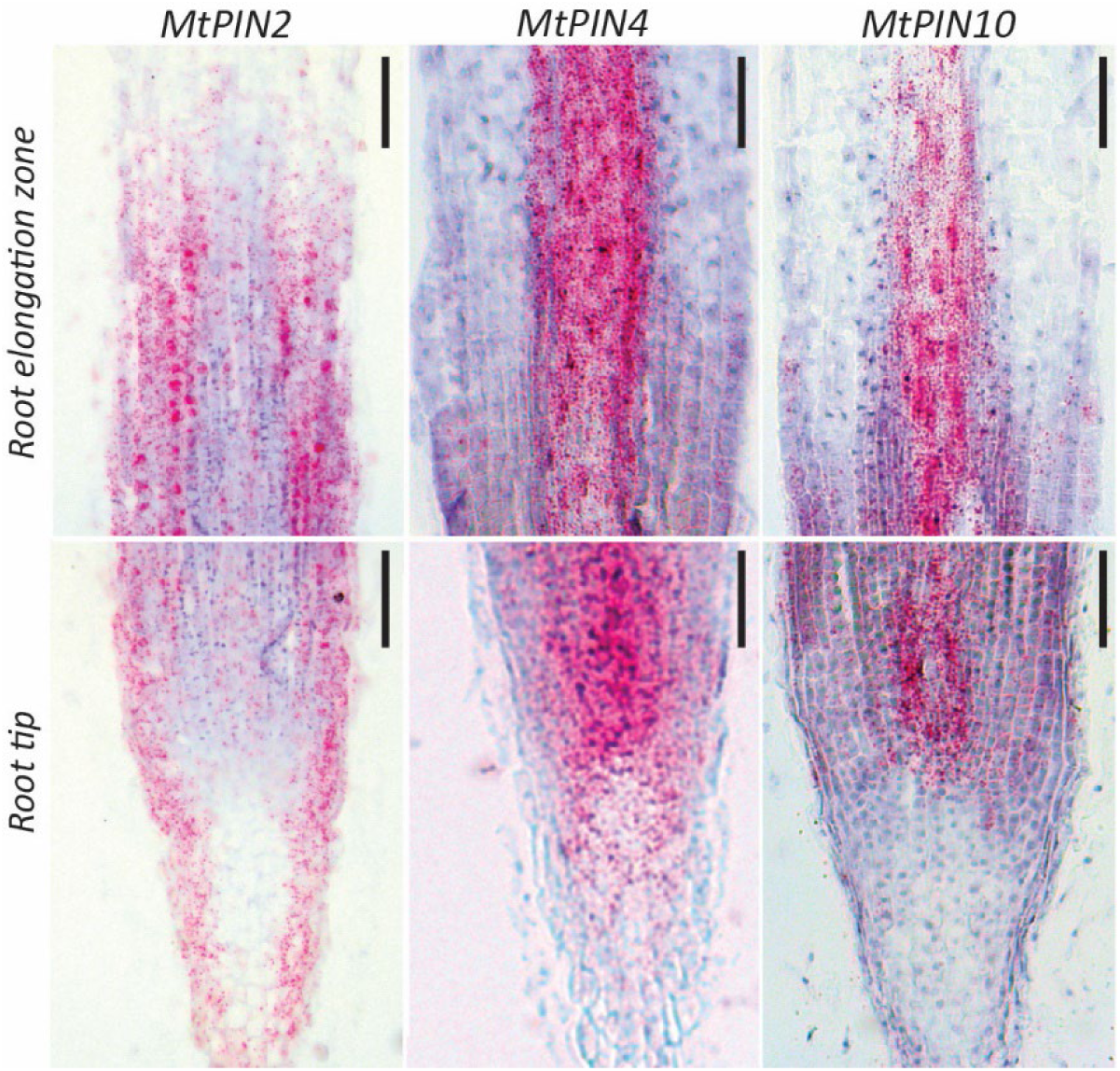
The spatiotemporal expression patterns of *PINs* in the root tip of Medicago A17. RNA *in situ* hybridizations with *MtPIN2, MtPIN4*, or *MtPIN10* probe set on longitudinal sections of the root tip and elongation zone (red dots are hybridization signals, scale bars 75 μm).

**Supplemental Figure S6:**
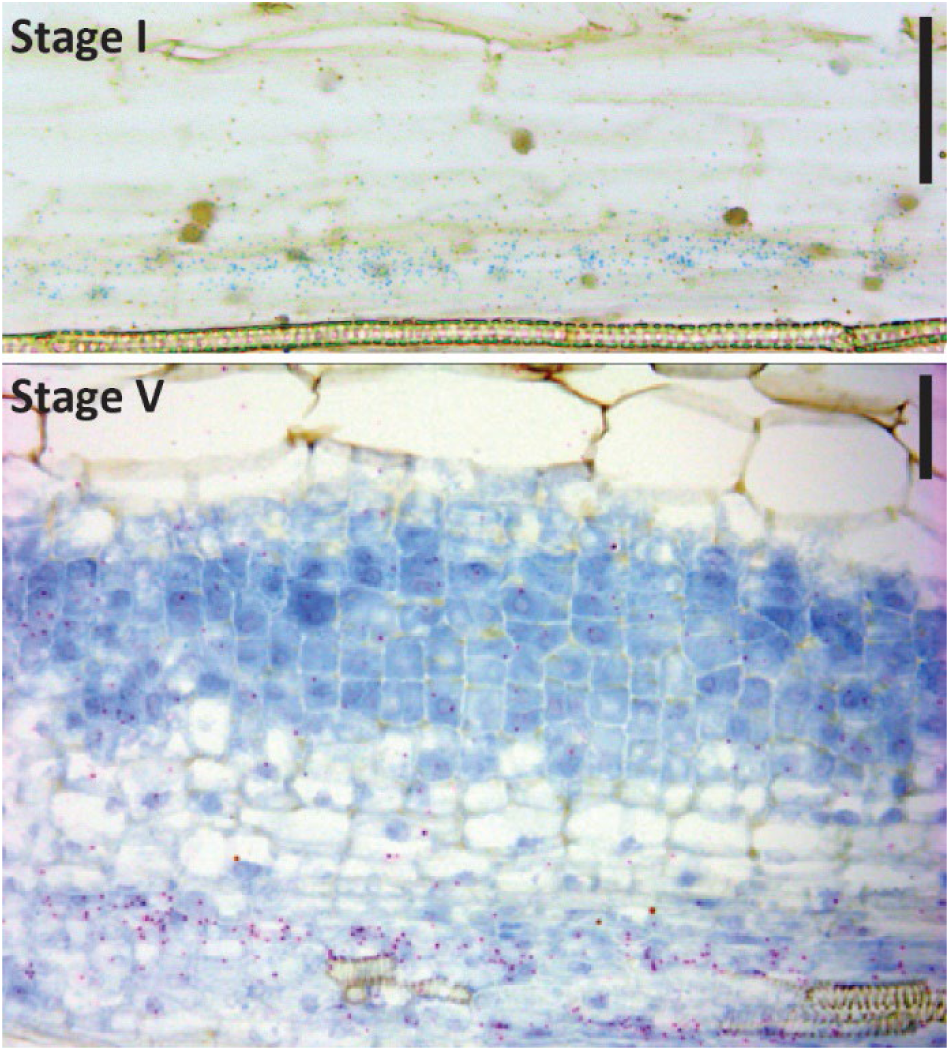
The spatiotemporal expression patterns of *MtPIN6* during nodule primordium formation. RNA *in situ* hybridization with *MtPIN6* probe sets on longitudinal sections of nodule primordia at stage I and V. For stage I, *NFYA1* was used as a marker for nodule primordium initiation. (Red dots are *MtPIN6* hybridization signals, blue dots are *MtNFYA1* hybridization signals, scale bars 75 μm).

**Supplemental Figure S7:**
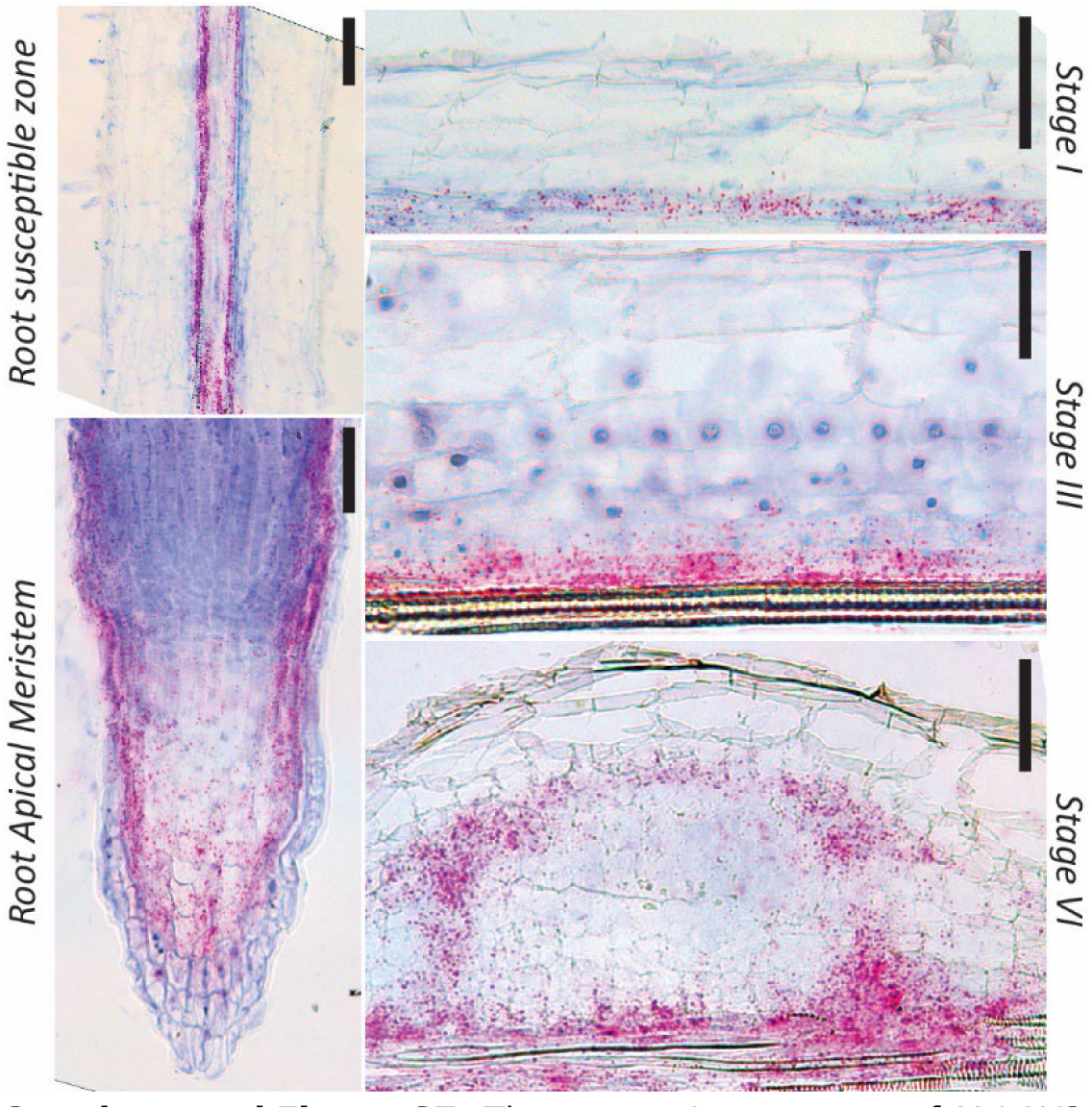
The expression patterns of *MtLAX2* in the root tip of, and during nodule primordium formation in Medicago A17. RNA *in situ* hybridization with *MtLAX2* probe set on longitudinal sections of the root susceptible zone, root tip and nodule primordia at stages I, III and VI (red dots are hybridization signals, scale bars 75 μm).

**Supplemental Figure S8.**
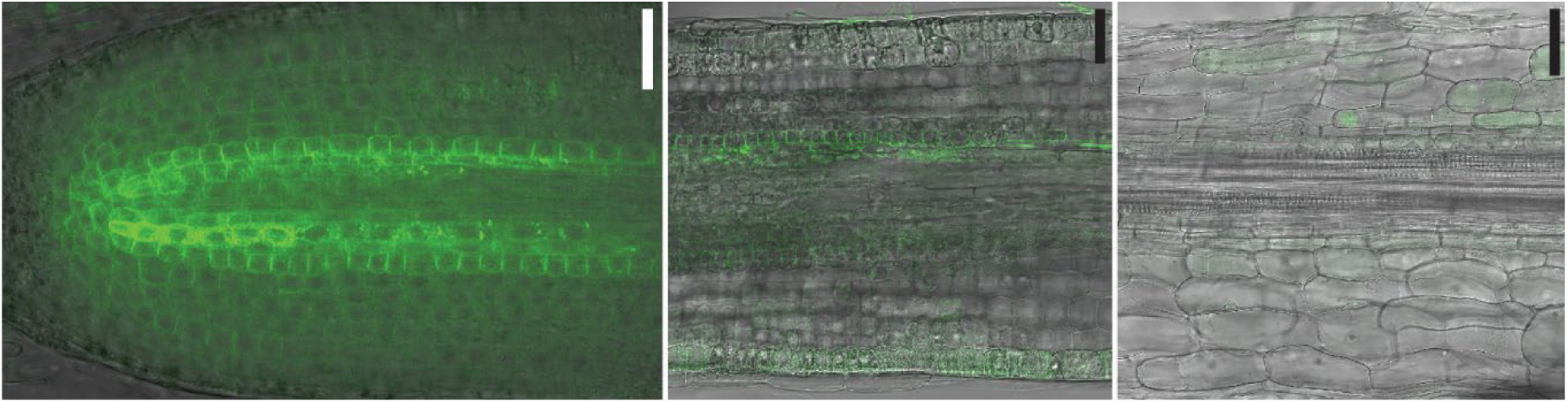
*Mt*PIN10 pattern in the Medicago root. *Mt*PIN10GFP is accumulated in vasculature and detected in cortical cells of root meristem, it is basal localized there. *Mt*PIN10GFP level decreased in elongation zone and become undetectable in root susceptible zone. (Sale bars 75 μm).

## SUPPLEMENTARY TABLES

**Table S1.**
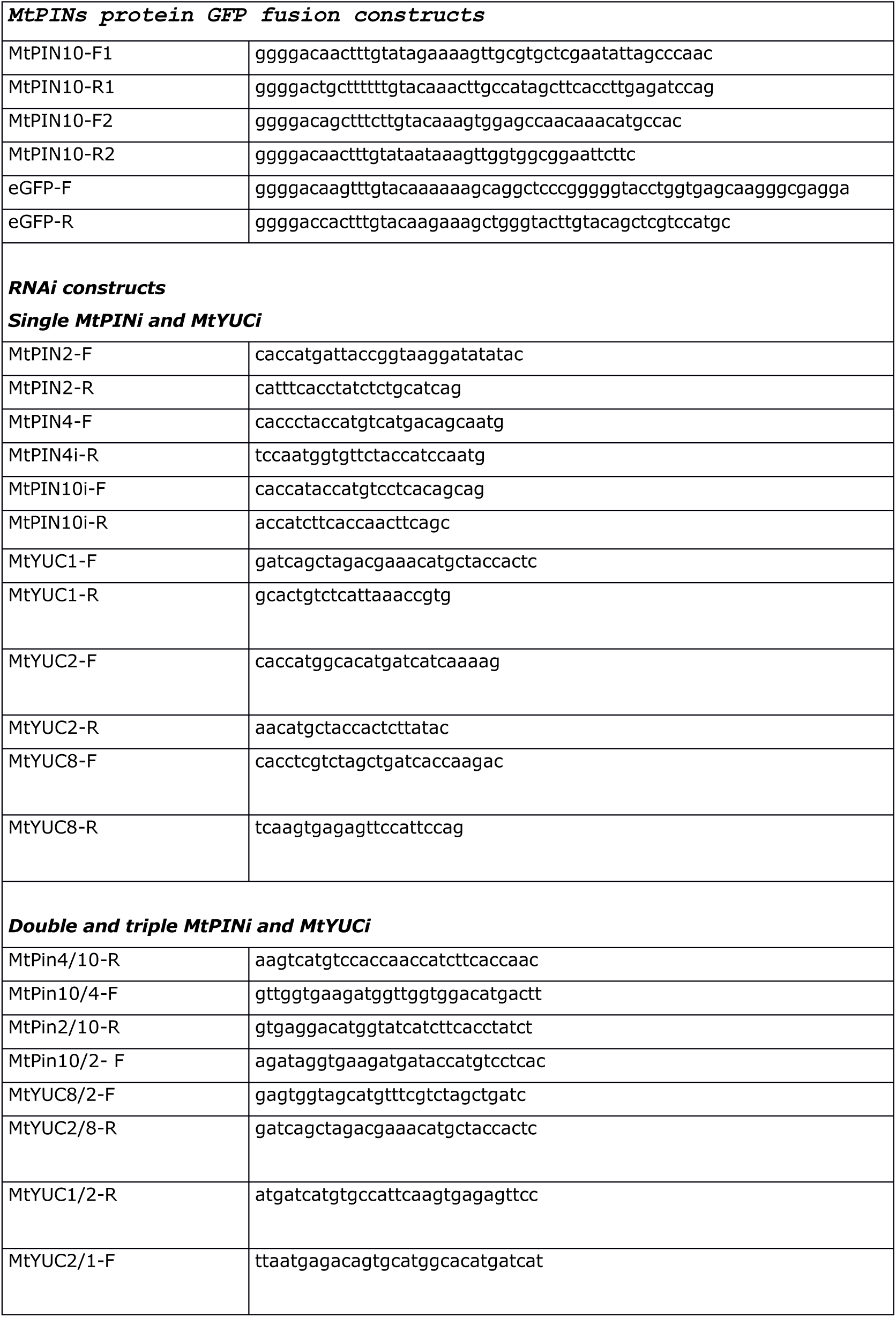
List of primers

**Table S2.** Gene probe sets used for ViewRNA *in situ* hybridization

| Gene name | Gene ID | Assay ID according to ThermoFisher Scientific |
| --- | --- | --- |
| *MtNF-YA1* | Medtr1g056530 | VPRWEK2 Type 6 |
| *MtYUC2* | Medtr6g086870 | VF1-6000670 Type 1 |
| *MtYUC8* | Medtr7g099330 | VF1-6000768 Type 1 |
| *MtYUC1* | Medtr3g109520 | VPKA3EK Type 1 |
| *MtYUC9* | Medtr1g069275 | VPCE3WM Type 1 |
| *MtPIN2* | Medtr4g127100 | VF1-17677 Type 1 |
| *MtPIN4* | Medtr6g069510 | VF1-20313 Type 1 |
| *MtPIN6* | Medtr1g029190 | VPDJXGJ Type 1 |
| *MtPIN10* | Medtr7g089360 | VF1-17681 Type 1 |
| *MtLAX2* | Medtr7g067450 | VPZTD33 Type 1 |
| *MtPIN9* | Medtr7g079720 | VF1-6000275 Type 1 |

